# LENG8 mediates RNA nuclear retention and degradation in eukaryotes

**DOI:** 10.1101/2025.08.14.670437

**Authors:** Lusong Tian, Liang Liu, Yoseop Yoon, Lindsey V. Soles, Marielle Valdez, Joshua Jeong, Clinton Yu, Lan Huang, Yongsheng Shi

**Affiliations:** Department of Microbiology and Molecular Genetics, School of Medicine, University of California, Irvine, Irvine, CA 92617, USA; Department of Physiology and Biophysics, University of California, Irvine, Irvine, CA 92697, USA; The Center for RNA Science and Therapeutics, University of California, Irvine, Irvine, CA 92697, USA

**Keywords:** RNA nuclear retention, intron retention, RNA export, RNA degradation, splicing, RNA polyadenylation

## Abstract

In eukaryotes, incompletely processed and misprocessed mRNAs as well as numerous noncoding RNAs are retained in the nucleus and often degraded. However, the mechanisms for this critical quality control pathway remain poorly understood. Here we identify LENG8 as a conserved RNA nuclear retention factor. We showed that LENG8 is recruited to pre-mRNAs by splicing factors, including the U1 snRNP. LENG8 binds to PCID2 and SEM1 to form the REX (Repressor of EXport) complex, which is conserved from yeast to human, and causes RNA nuclear retention by acting as a dominant negative factor for the essential mRNA export factor TREX-2. LENG8 depletion leads to the leakage of misprocessed mRNAs, including intronically polyadenylated and intron-retained mRNAs, as well as noncoding RNAs into the cytoplasm. Finally, LENG8 promotes RNA degradation by recruiting PAXT and the RNA exosome. Thus our study revealed a conserved quality control mechanism for eukaryotic gene expression that ensures only fully and correctly processed RNAs are exported from the nucleus.

## Introduction

In eukaryotic cells, the separation of mRNA biogenesis and translation into distinct compartments allows for exquisite quality control. A key aspect of this surveillance mechanism is to ensure only correctly processed RNAs are exported from the nucleus. Typical pre-mRNAs undergo multiple processing steps, including 5′ capping, splicing, and cleavage and polyadenylation (CPA)^1,2^. The TREX (Transcription-Export) complex plays an essential role in coupling mRNA processing with export, and it consists of the THO subcomplex, the RNA helicase UAP56 (DDX39B), and mRNA export adaptor proteins including Aly/REF^3,4^. TREX can be recruited to the 5′ end, splice junctions, and the 3′ end of mRNAs via interactions with the cap-binding complex, the exon junction complex and the CPA machinery^5–7^. In turn, the TREX complex helps to recruit the mRNA export receptor NXF1-NXT1 complex to assemble an export competent mRNA-protein complex^3,4,8^. Subsequently mRNAs are transferred to the nuclear pore complex (NPC)-bound TREX-2 complex for export^3,4,9^.

To safeguard gene expression fidelity, eukaryotes have evolved mechanisms to retain incompletely processed mRNAs, such as intron-retained RNAs, in the nucleus^10,11^. Additionally, many long noncoding RNAs are naturally retained in the nucleus^12,13^. Importantly, misprocessed mRNAs, including intronically polyadenylated mRNAs (IPAs), are retained and degraded in the nucleus^14–16^. Several factors have been implicated in RNA nuclear retention, including the yeast protein Mpl1p and its mammalian homologue TPR, early spliceosome components and the RNA exosome adaptor PAXT^15,17–19^. However, it is unclear whether their putative function in RNA nuclear retention is direct or indirect. Two long-standing models have been proposed for RNA nuclear retention. The first model posits that RNA nuclear retention is due to defective recruitment of RNA export factors. As the TREX complex is recruited to RNAs in an RNA processing-dependent manner, defective or inefficient processing could lead to decreased loading of RNA export factors and, consequently, nuclear retention^10,20^. It was believed that many long noncoding RNAs are retained in the nuclear due to their inefficient splicing. However, TREX complex can be recruited to the 5′ end or the 3′ end in a splicing-independent manner^6,7,21^. Additionally, transcripts of multi-intron genes with only one unspliced intron can be sequestered in the nucleus despite the fact that multiple EJCs have been deposited on the same RNA molecule^22^. Therefore, RNA nuclear retention cannot be explained solely by defective loading of export factor. On the other hand, 5’ splice site (ss) and U1 snRNP have been shown to retain RNAs in the nucleus from yeast to human, presumably by recruiting putative nuclear retention factors^23–25^. However, no such nuclear retention factor has been identified.

In this study, we identified LENG8, a poorly characterized protein that is conserved from yeast to human, as a key nuclear retention factor. It is recruited to RNAs via its association with early splicing factors, including U1 snRNP. LENG8 forms an evolutionarily conserved complex called REX and blocks RNA export by acting as a dominant negative factor for TREX-2. Finally, LENG8 is required for recruiting PAXT and the RNA exosome for degrading mis-processed mRNAs and noncoding RNAs.

## Results

### Identify novel factors involved in 5′ ss-dependent RNA nuclear retention and degradation using multiple unbiased approaches

We have recently shown that RNAs that contain the combination of a 5’ ss and a poly(A) junction, such as intronically polyadenylated mRNAs, are retained in the nucleus and targeted for degradation by the RNA exosome and its adaptor PAXT (ZFC3H1/MTR4)^14^. To better understand the mechanism for 5’ ss-dependent RNA nuclear retention and degradation, we first performed a genome-wide CRISPR knockout screen. To this end, we constructed eGFP-5′ ss-poly(A) site (PAS) reporter, in which a wild-type (WT) or mutant (Mut) 5′ss was inserted between the eGFP coding sequence and a PAS and the reporter expression was driven by a Tet-inducible promoter (Figure **1A**). These reporters were stably transfected into the Flp-In 293 cell line and, consistent with our previous study^14^, the eGFP signals from cells expressing WT 5′ ss-containing reporter were significantly lower than those from cells expressing the Mut 5′ ss-containing reporter (Figure **S1A**). Furthermore, treatment of cells expressing WT 5′ss-containing reporter with a U1 antisense morpholino oligo (AMO), which blocks U1-RNA interaction, led to drastic increase in eGFP signals (Figure **S1B**). We have shown previously that this reporter can faithfully recapitulate 5′ ss-dependent mRNA nuclear retention and degradation and thus we used this cell line to identify novel factors involved in this pathway.

**Figure 1.**
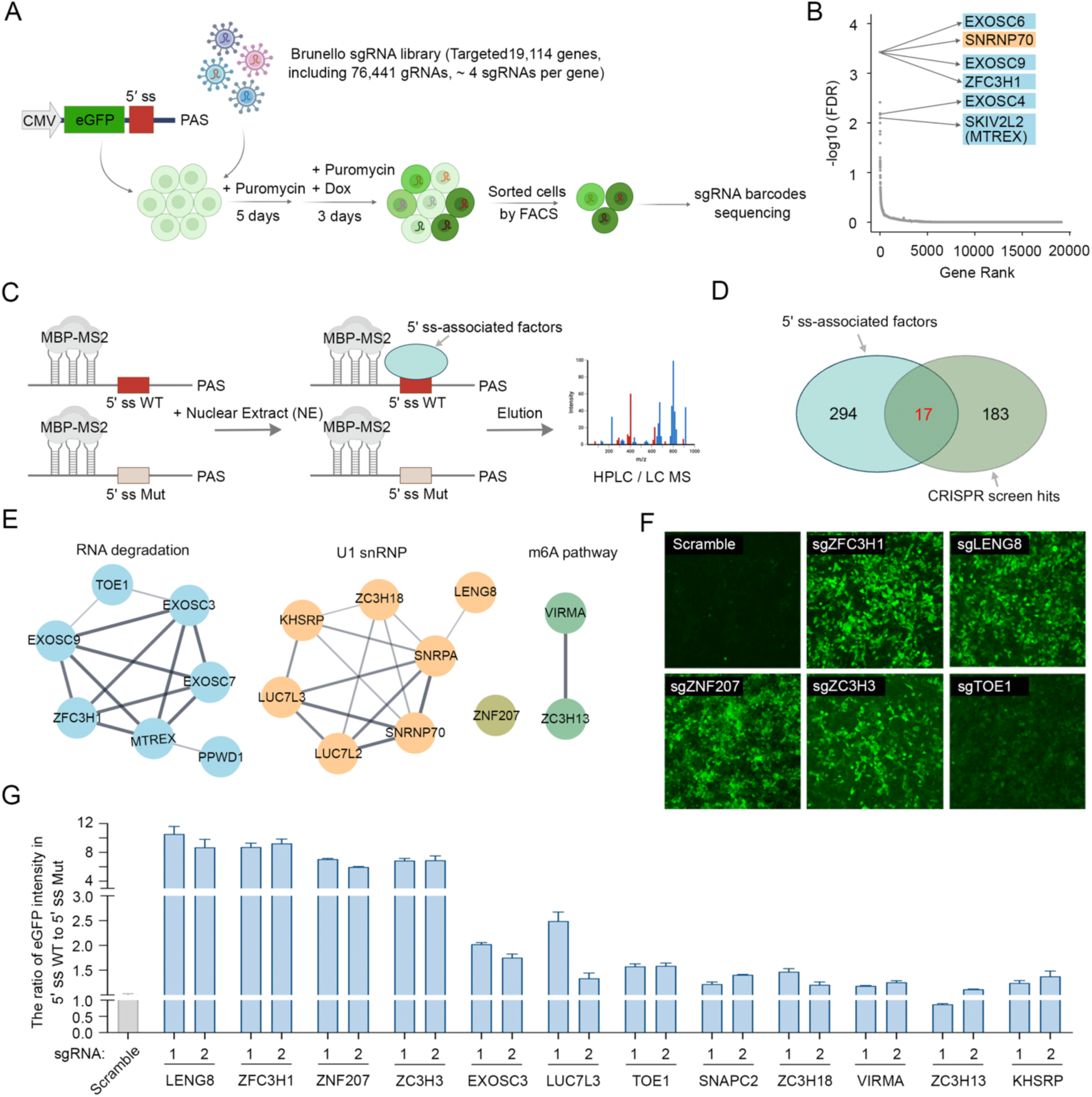
Identify novel factors involved in 5′ ss-dependent RNA nuclear retention and degradation using multiple unbiased approaches. (A) Workflow of a genome-wide CRISPR screen to identify candidate factors involved in 5′ ss-mediated gene repression. 293 Flp-In cells expressing the eGFP-5′ ss-PAS reporter are infected with a lentiviral Brunello genome-wide sgRNA library and selected with puromycin for 5 days. The selected cells are then treated with doxycycline for 3 days to induce eGFP expression. Following induction, the top 1% of high eGFP-expressing cells are sorted via FACS analysis. Genomic DNA is then extracted, followed by sgRNA barcode amplification and deep sequencing analysis. (B) Candidate genes identified by the CRISPR-Cas9 screen. Data analysis is performed using the MAGeCK algorithm to identify enriched sgRNA, and the genes are ranked by false discovery rate (FDR). Names of top candidates were labeled. (C) A schematic diagram for the RNA pull-down and mass spectrometry analysis. MBP-MS2, a fusion protein combining maltose-binding protein (MBP) and MS2, facilitating RNA capture and purification. 5′ ss: 5′ splicing site; WT: wild type; Mut: mutant. (D) A Venn diagram showing the overlap between the 5′ ss-associated factors and hits from our CRISPR screen. (E) STRING network analysis of the 17 shared factors. (F) eGFP signals from the eGFP-5′ ss-PAS reporter cell line treated with scramble or specific sgRNAs against the specified factors (related to **Extended Data Fig. 1H**). (G) Quantified eGFP signals from the eGFP-5′ ss-PAS reporter cell line treated with scramble or specific sgRNAs against the specified factors.

For the screen, the eGFP-WT 5′ ss-PAS reporter cell line was infected with the Brunello lentiviral sgRNA library (Figure **1A**), which targets 19,114 genes with a total of 77,441 sgRNAs (4 sgRNAs per gene)^26^. Following selection for sgRNA-expressing cells, reporter expression was induced by adding Doxycycline (Figure **1A**). As depletion of any factors required for 5′ ss-dependent mRNA nuclear retention or degradation was expected to increase eGFP signals, we selected cells with the top 1% eGFP signals using fluorescence-activated cell sorting (FACS), extracted genomic DNA from these cells, and identified the sgRNA barcodes contained in these cells by sequencing (Figure **1A** and Figure **S1C**). The top 200 candidate genes from the CRISPR screen with the lowest false discovery rate included the RNA exosome subunits, the exosome adaptor PAXT (ZFC3H1 and MTREX) and U1 snRNP components (Figure **1B**, Figure **S1D, S1E**, and **Supplemental Table 1**), fully consistent with our previously proposed degradation model^14^. Additionally nuclear speckle components and m6A methyltransferase complex components were also enriched (Figure **S1D, S1E**, and **Supplemental Table 1**), consistent with previous reports implicating these factors in RNA degradation or retention^27,28^. These findings further validated the robustness and reliability of our CRISPR screen in uncovering novel factors required for 5′ ss-dependent RNA nuclear retention and degradation.

The factors identified in our CRISPR screen could influence 5′ ss-dependent RNA nuclear retention and/or degradation directly or indirectly. To identify the factors that directly participate in these processes, we hypothesized that they must be recruited to 5′ ss-containing RNAs. To identify factors that bind to RNA in a 5′ ss-dependent manner, we performed RNA pulldown assays using RNAs containing WT or Mut 5′ ss fused with three copies of the MS2 hairpin (Figure **1C**). We first incubated the RNAs with MBP (maltose binding protein)-MS2 fusion protein followed by incubation with HeLa cell nuclear extract. The assembled RNA-protein complexes were pulled down using amylose beads and the proteins that specifically associated with the WT 5′ ss were identified by mass spectrometry (MS) analyses (Figure **1C**). 311 proteins are identified as 5′ ss-associated factors (Fold Enrichment WT/Mut 5′ ss ≥ 1.5, **Supplemental Table 2**) and, as expected, they contained the U1 snRNP and other splicing factors (Figure **S1F, S1G**, **Supplemental Table 3**). A comparison of these 5′ ss-associated proteins with our CRISPR screen hits identified 17 shared factors (Figure **1D**), which contain U1 snRNP and associated proteins, RNA degradation factors, m6A modification factors, and other less characterized proteins (Figure **1E**). Consistent with our results, ZNF207, a known cell cycle regulator, has recently been identified as a regulator of splicing via interactions with U1 snRNP ^29^. To validate the functional roles of these factors, all 17 shared factors were depleted using CRISPR/Cas9 in cells stably expressing the eGFP-5′ ss WT or Mut reporters and we observed significant increases in eGFP signals in a 5′ ss-dependent manner, particularly upon depletion of ZFC3H1, LENG8, ZC3H3, and ZNF207 (Figure **1F**, **1G**, and Figure **S1H**). Together, by using two unbiased approaches, we have identified a set of factors that are associated with 5′ ss and repressed reporter gene expression in a 5′ ss-dependent manner.

### LENG8 mediates 5′ ss-dependent nuclear retention and degradation of RNAs

We have shown previously that the expression of the 5′ ss-PAS-containing reporter is repressed by nuclear retention and degradation^14^. To determine the precise functions our candidate factors in these processes, we depleted them in cells expressing the eGFP-5′ ss reporter and measured the reporter mRNA levels by qRT-PCR. For the top four candidate genes, we generated HEK293T cell lines in which a FKBP12^F36V^ degron was fused to the endogenous genes by using CRISPR/Cas9^30,31^. Treatment of these edited cell lines with a small molecule, dTAG-V1, resulted in near complete depletion of the candidate proteins via the ubiquitin-proteasome system (Figure **S2A**). Next, we examined the expression of the eGFP-5′ ss WT-PAS or eGFP-5′ ss Mut-PAS reporter expression in DMSO- or dTAG-V1-treated cells (Figure **2A**). The results showed that eGFP intensity as well as the reporter mRNA levels significantly increased in cells expressing eGFP-5′ ss WT-PAS upon depletion of ZFC3H1, ZNF207, LENG8 and ZC3H3, while minimal changes were observed in cells expressing eGFP-5′ ss Mut-PAS (Figure **2B** and Figure **S2B**). These results suggest that ZFC3H1, ZNF207, LENG8, and ZC3H3 are all required for 5′ ss-dependent RNA degradation.

**Figure 2.**
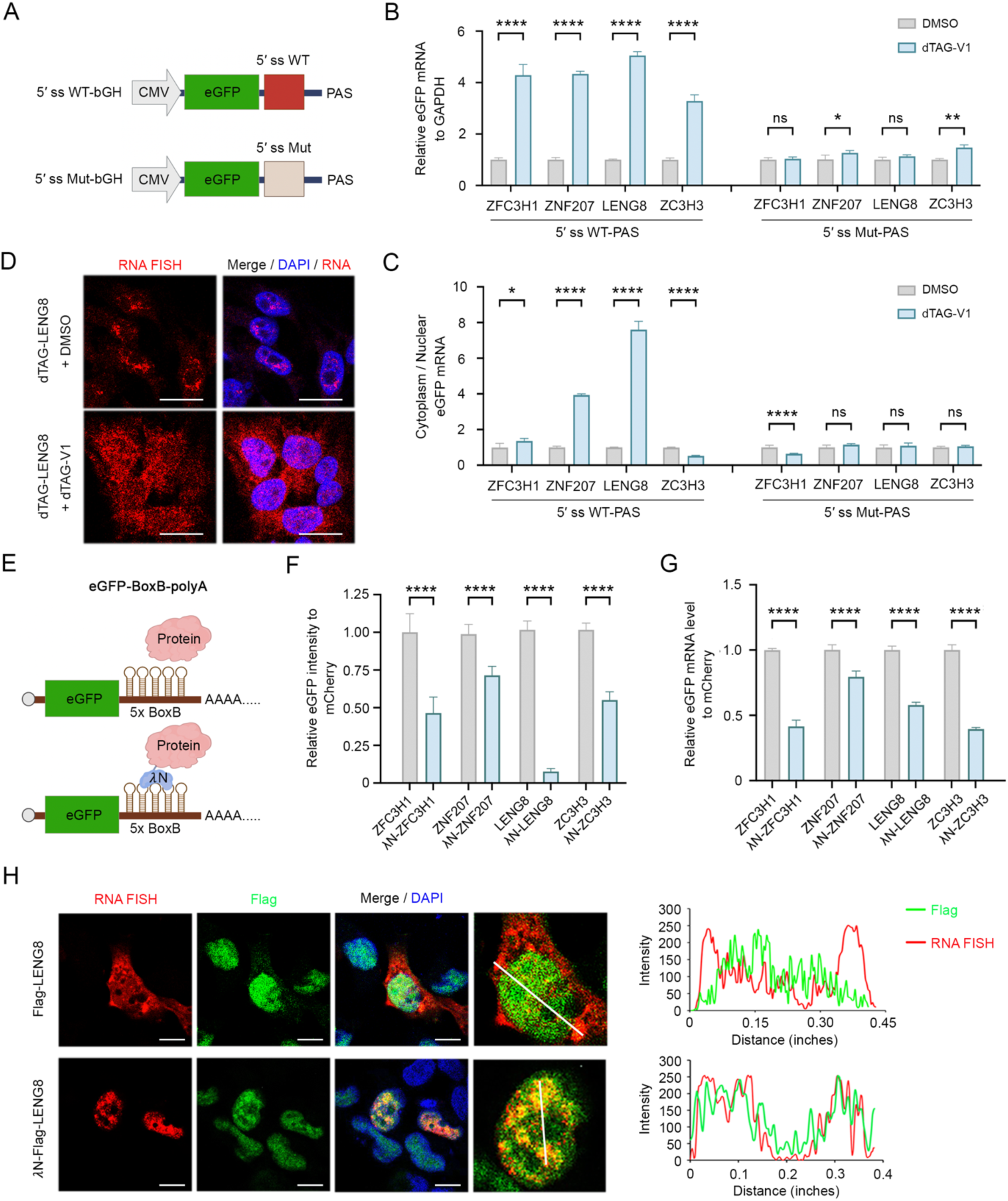
LENG8 mediates 5′ ss-dependent nuclear retention and degradation of RNAs. (A) Schematic of the eGFP-5′ ss-PAS reporter containing a WT or Mut 5′ ss in the 3′ untranslated region. (B) Reporter mRNA level is measured in cells expressing either the eGFP-5′ ss WT-PAS (left group) 5′ ss Mut (right group) following depletion of ZFC3H1, ZNF207, LENG8 and ZC3H3, respectively. The relative eGFP mRNA level is normalized to GAPDH and then quantified in comparison to the DMSO condition. Data presented as mean ± SEM (n = 3). *p value % 0.05, **p value % 0.01, ***p value % 0.001, ****: p-value % 0.0001, ns: not significant (unpaired t-test). (C) The relative cytoplasmic eGFP mRNA level is measured in cells expressing reporters following the depletion of ZFC3H1, ZNF207, LENG8 and ZC3H3, respectively. Cytoplasmic eGFP mRNA level is normalized to nuclear mRNA and quantified relative to the DMSO control condition. Data presented as mean ± SEM (n = 3). *p value % 0.05, **p value % 0.01, ***p value % 0.001, ****: p-value % 0.0001, ns: not significant (unpaired t-test). (D) RNA FISH analysis of eGFP-5′ ss WT-PAS reporter is performed in DMSO- or dTAG-V1-treated LENG8-FKBP12^F36V^ cell line. The reporter mRNA signal is shown in red, while nuclei are stained with DAPI and depicted in blue. Scale bars: 20 μm. (E) Schematic of the eGFP-BoxB-PAS reporter. These constructs contain the eGFP coding sequence followed by a 5×BoxB motif, which can be bound by the λN peptide. (F) eGFP signals in cells expressing candidate proteins without or with a λN peptide fusion. The signals in cells expressing candidate proteins are normalized to those of a co-transfected mCherry. Data presented as mean ± SEM (n = 3). *p value % 0.05, **p value % 0.01, ***p value % 0.001, ****: p-value % 0.0001 (unpaired t-test). (G) eGFP mRNA level is measured in cells co-expressing the eGFP-BoxB-polyA reporter and candidate proteins, either with or without an N-terminal λN peptide. The relative eGFP mRNA level is normalized to that of a co-transfected mCherry control plasmid and then quantified relative to the no λN peptide control condition. Data presented as mean ± SEM (n = 3). *p value % 0.05, **p value % 0.01, ***p value % 0.001, ****: p-value % 0.0001 (unpaired t-test). (H) Reporter RNA FISH and FLAG immunofluorescence (IF) analysis in stable Flp-In 293 cells expressing either FLAG-LENG8 or λN-FLAG-LENG8. The reporter schematic is showed in Extended Data 2j. The RNA signal from eGFP-BoxB-MALAT1 reporter is shown in red, while the FLAG-LENG8 IF signal is shown in green. Nuclei is stained with DAPI (blue). The graphs in the right panel display the intensity profiles of RNA FISH and FLAG signals along the straight line drawn in the middle panel. Scale bars: 20 μm.

We next investigated the functions of these factors in RNA nuclear retention by fractionating DMSO- or dTAG V1-treated the degron cell lines into nuclear and cytoplasmic fractions, extracting RNAs, and comparing the reporter mRNA levels in these compartments by qRT-PCR (Figure **S2C**). The results showed that the cytoplasmic/nuclear ratio of the eGFP-5′ ss WT-PAS reporter mRNAs increased by ∼4- and ∼8-fold following depletion of ZNF207 or LENG8 respectively (Figure **2C**). A modest increase was also observed for ZFC3H1-depleted cells while ZC3H3 depletion caused a modest decrease in cytoplasmic/nuclear RNA ratio (Figure **2C**). Little or no change was observed for the eGFP-5′ ss Mut-PAS reporter (Figure **2C**). These results suggest that LENG8 and, to a lesser degree, ZNF207 are required for 5′ ss-dependent RNA nuclear retention. To directly visualize RNA localization, we performed RNA fluorescent in situ hybridization (FISH) analysis of the eGFP-5′ ss WT-PAS reporter mRNAs in control or ZFC3H1-, ZNF207-, LENG8- and ZC3H3-depleted cells (Figure **2D**, Figure **S2D**). Consistent with the qRT-PCR results (Figure **2B**), we observed significantly higher overall RNA FISH signals in cells depleted of these factors (Figure **2D** and Figure **S2D**). Interestingly, the higher RNA FISH signals were concentrated in the nuclei of ZFC3H1 or ZC3H3-depleted cells (Figure S**2D**). In contrast, higher RNA FISH signals were observed in both the nucleus and cytoplasm of LENG8-depleted cells (Figure **2D**), suggesting that LENG8 is required for nuclear retention of the reporter mRNAs. Depletion of ZNF207 also led to significant spread of reporter mRNA into the cytoplasm (Figure **S2D**). Of note, we observed more mitotic cells upon ZNF207 depletion (Figure **S2D**), consistent with its known function in mitosis^32–34^. Together these results suggest that, although all four factors are required for 5′ ss-dependent RNA degradation, LENG8 and ZNF207 are necessary for 5′ ss-dependent RNA nuclear retention.

In the eGFP-5′ ss-PAS reporter, RNA nuclear retention and degradation was tightly coupled. In our recent study, we showed that if the PAS was replaced by the *MALAT1* 3’ end sequence, which forms a nonpolyadenylated triple-helix structure^35^, the reporter was retained in the nucleus in a 5′ ss-dependent manner, but not degraded^3^. To directly test the functions of our candidate factors in nuclear retention, we tested the effect of depleting these factors on these the eGFP-5′ ss-MALAT1 reporter expression (Figure **S2E**). Upon depletion of ZNF207, LENG8, and ZC3H3, but not ZFC3H1, the eGFP signal significantly increased, accompanied by modest increases in mRNA levels (Figure **S2F, S2G**). When we examined the cytoplasmic/nuclear mRNA ratio, LENG8 depletion led to the highest increase while depleting other factors had minimal effect (Figure **S2H**), providing further evidence that LENG8 is necessary for 5′ ss-dependent nuclear retention.

To test whether the candidate factors are sufficient for RNA degradation and nuclear retention, a tethering assay was performed using a reporter in which the eGFP coding sequence is followed by a 5×BoxB motif (Figure **2E**). This motif can be bound by the λN peptide, allowing for targeted tethering of specific proteins to the reporter RNA^36^. Using this strategy, we tethered FLAG-tagged ZFC3H1, ZNF207, LENG8 and ZC3H3 to the eGFP-5×BoxB reporter and measured the eGFP signal and reporter mRNA levels. The results showed that tethering any of the four factors to the reporter mRNA significantly decreased eGFP intensity, with LENG8 exhibiting the strongest effect Figure **2F**). eGFP mRNA levels also decreased upon tethering of these factors (Figure **2G**), indicating that RNA degradation was induced. Importantly LENG8 tethering caused over 11-fold decrease in eGFP signals, but less than 2-fold decrease in mRNA levels (compare Figure 2F and 2G), suggesting that LENG8 tethering could cause a defect in reporter mRNA export. Indeed, RNA FISH combined with FLAG immunofluorescence analysis revealed that, although untethered reporter mRNAs were found in both nucleus and cytoplasm, tethering LENG8 to the reporter mRNAs restricted them in the nucleus (Figure **2H**). Together, these analyses showed that ZFC3H1, ZNF207, LENG8, and ZC3H3 are all involved in the degradation of 5′ ss-containing RNAs. Most importantly, our results demonstrated that LENG8 is necessary and sufficient for retaining 5′ ss-containing RNAs in the nucleus.

### LENG8 mediates nuclear retention and degradation of misprocessed mRNAs and noncoding RNAs at the transcriptome-wide level

Our reporter-based results suggest that LENG8 mediates nuclear retention of 5′ ss-containing RNA and the degradation of RNAs containing a combination of 5′ ss and poly(A) junction (Figure 2). We next wanted to determine the impact of LENG8 on the stability and subcellular localization of endogenous RNAs. To this end, we treated the LENG8-FKBP12^F36V^ degron cell line with DMSO or dTAG-V1, extracted RNAs from whole cells or the nuclear and cytoplasmic fractions and subjected them to mRNA-seq and PAS-seq analyses^37^, the latter maps the poly(A) junctions of RNAs and provides quantitative measurement of poly(A) site usage and transcript abundance. Our previous study demonstrated that IPAs were the major class of misprocessed RNAs for PAXT-mediated exosomal RNA degradation because they contain the combination of 5′ ss and poly(A) junction^14^. Similarly, we found that LENG8 depletion led to significant accumulation of IPAs while non-IPA mRNA levels did not change (Figure **3A**, Figure **S3A**). Importantly, we observed more significant increase in cytoplasmic IPA levels compared to that in the nucleus (Figure **3B**, Figure **S3B, S3C, S3D**), suggesting that LENG8 depletion led to both accumulation and export of IPAs. This trend was exemplified by two example genes, *PLPBP* and *GAA*. As shown in Figure 3C (left panel), the IPA isoform of *PLPBP* was restricted to the nucleus in DMSO-treated cells but was released into the cytoplasm in dTAG-V1 treated cells. Low level of IPA isoform of GAA was observed in the nucleus of control cells, but it accumulated to much higher levels and was exported into the cytoplasm in LENG8-depleted cells (Figure **3C**, right panel), suggesting that LENG8 mediates both nuclear retention and degradation of IPA RNAs.

**Figure 3.**
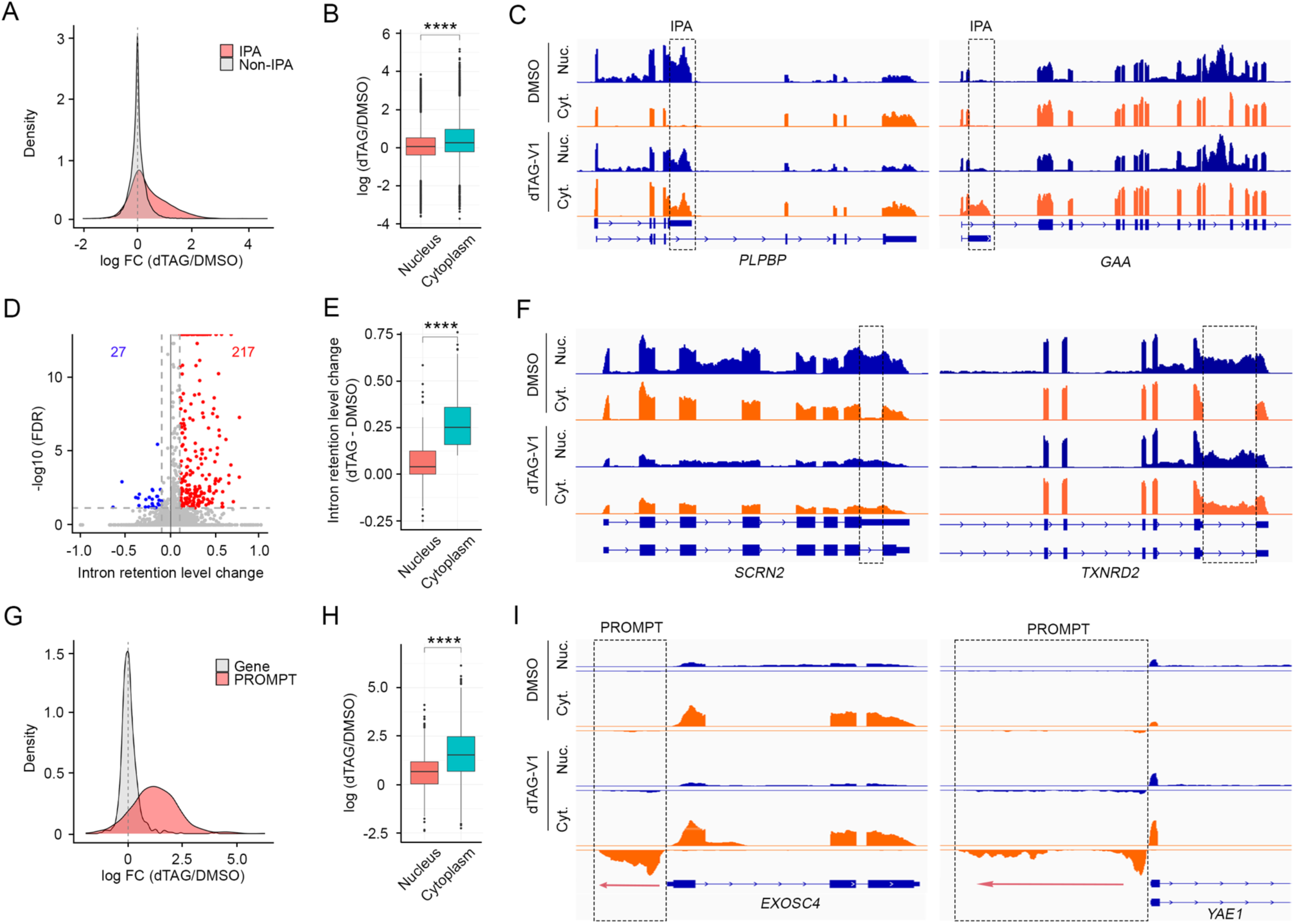
LENG8 mediates nuclear retention and degradation of misprocessed mRNAs and noncoding RNAs at the transcriptome-wide level. (A) Density plots of the log_2_FoldChange (log_2_FC) for all IPA or non-IPA transcripts following depletion of LENG8 in HEK293T cells. IPA transcripts are depicted in red and non-IPA transcripts grey. (B) Box plot showing the relative IPA levels in dTAG- vs. DMSO-treated cells (log2(dTAG/DMSO)). Statistical analysis is calculated by Mann-Whitney test. ****p value ≤ 0.0001. (C) mRNA-seq tracks for the genes *PLPBP* and *GAA* in cytoplasm and nucleus in DMSO- and dTAG-V1-treatment LENG8-FKBP12^F36V^ cells. Regions unique to the IPA transcripts are marked by boxes. (D) A volcano plot showing intron inclusion level change in the cytoplasm after depletion of LENG8. (E) Box plot depicting that the relative levels of intron-retained transcripts in the nucleus and cytoplasm of dTAG vs DMSO-treated LENG8-FKBP12^F36V^ cells. Statistical analysis is calculated by Mann-Whitney test. ****p value ≤ 0.0001. (F) mRNA-seq racks for the genes *SCRN2* and *TXNRD2* in cytoplasm and nucleus in DMSO- and dTAG-V1-treated LENG8-FKBP12^F36V^ cells. (G) Density plots of the log_2_FoldChange (log_2_FC) for all PROMPT and the corresponding protein-coding transcripts following depletion of LENG8 in HEK293T cells. PROMPT transcripts are depicted in red and normal gene transcripts grey. (H) Box plot depicting the relative levels of PROMPTs in the nucleus and cytoplasm of dTAG vs DMSO-treated LENG8-FKBP12^F36V^ cells. Statistical analysis is calculated by Mann-Whitney test. ****p value ≤ 0.0001. (I) mRNA-seq data tracks for the genes *EXOSC4* and *YAE1* in cytoplasm and nuclei in DMSO- and dTAG-V1-treated LENG8-FKBP12^F36V^ cells.

In addition to IPAs, we also observed significant accumulation of intron-retained mRNAs in the cytoplasm of LENG8-depleted cells (Figure **3D**). Additionally, the intron retention level for these RNAs was significantly higher in the cytoplasm compared to that in the nucleus (Figure **3E**), suggesting that LENG8 depletion led to the export of these intron-retained mRNAs. As shown in Figure 3F, intron-retained mRNAs of *SCRN2* and *TXNRD2* genes were restricted to the nucleus in control cells but were released into the cytoplasm in LENG8-depleted cells. Interestingly, the abnormally exported intron-retained mRNAs in LENG8-depleted cells were highly enriched for retained last introns while the first or middle introns were depleted (Figure **S3E**). The mechanistic implications for this observation will be discussed later.

The degradation and nuclear retention defects in LENG8-depleted cells were not limited to mRNAs as we also observed the accumulation and export of many noncoding RNAs. One prominent group is PROMPTs (promoter upstream antisense transcripts), which are known to be highly unstable due to RNA exosome-mediated degradation^38^. In LENG8-depleted cells, PROMPTs levels drastically increased while the corresponding transcripts of the protein-coding genes from the same promoters did not change (Figure **3G**). Similar to IPAs and intron-retained mRNAs, the increase in PROMPT levels was significantly higher in the cytoplasm than in the nucleus (Figure **3H**, Figure **S3F, S3G, S3H**), suggesting that LENG8 depletion led to defects in both the degradation and nuclear retention of these transcripts, as shown by two examples (Figure **3I**). Taken together, these results strongly suggest that LENG8 broadly mediates the nuclear retention of intron-retained mRNAs, IPA transcripts, and noncoding RNAs, as well as the degradation of IPAs and noncoding RNAs.

### LENG8 is recruited to target RNAs via interactions with U1 snRNP and other splicing factors

To investigate how LENG8 is recruited to its target RNAs, we first analyzed its protein interactome. To this end, we over-expressed FLAG-tagged LENG8 and performed immunoprecipitation (IP) using anti-FLAG antibody followed by MS analysis. Gene ontology (GO) analysis of LENG8-associated factors (Fold Enrichment ≥ 1.5 in both replicates, **Supplemental Table 4**) showed significant enrichment of pathways related to nucleocytoplasmic transport, the spliceosome, and the RNA surveillance pathway (Figure **4A**, **Supplemental Table 5**), consistent with LENG8’s role in RNA nuclear retention and degradation. The splicing factors associated with LENG8 included all the major splicing snRNPs (Figure **4B**). As our reporter assays and genome-wide analyses showed that LENG8 specifically target 5′ ss-containing RNAs and 5′ ss is recognized by U1 snRNP, we next analyzed the potential interactions between LENG8 and U1 snRNP. First, we performed FLAG IP of full-length (FL) LENG8 and confirmed that U1 snRNP components were co-precipitated (Figure **S4A**). Next, we examined the role of different LENG8 regions in mediating interactions with U1 snRNP. LENG8 protein contains a long disordered region (1-540 amino acid (aa)) at its N-terminus and a C-terminal SAC3/GANP/THP3 domain (541-800aa, referred to SAC3 domain herein) (Figure **4C, 4D**). When we expressed FLAG-tagged N- or C-terminal regions of LENG8 and performed FLAG IP, we found that the N-terminal region interacted strongly with U1 snRNP while the C-terminal region did not (Figure **S4A**). Although the N-terminal region of LENG8 is largely disordered, a structured region was predicted for the middle portion (290-330aa) (Figure **4C, 4D**). We next divided the N-terminal region (1-540aa) into three segments: 1-290aa, 200-400aa, and 330-540aa (Figure **4D**). Intriguingly, we observed that both 1-290aa and 330-540aa regions interacted with U1 snRNP in an RNA-independent manner (Figure **4E**), suggesting that both regions of LENG8 interact with one or more protein components of the U1 snRNP.

**Figure 4.**
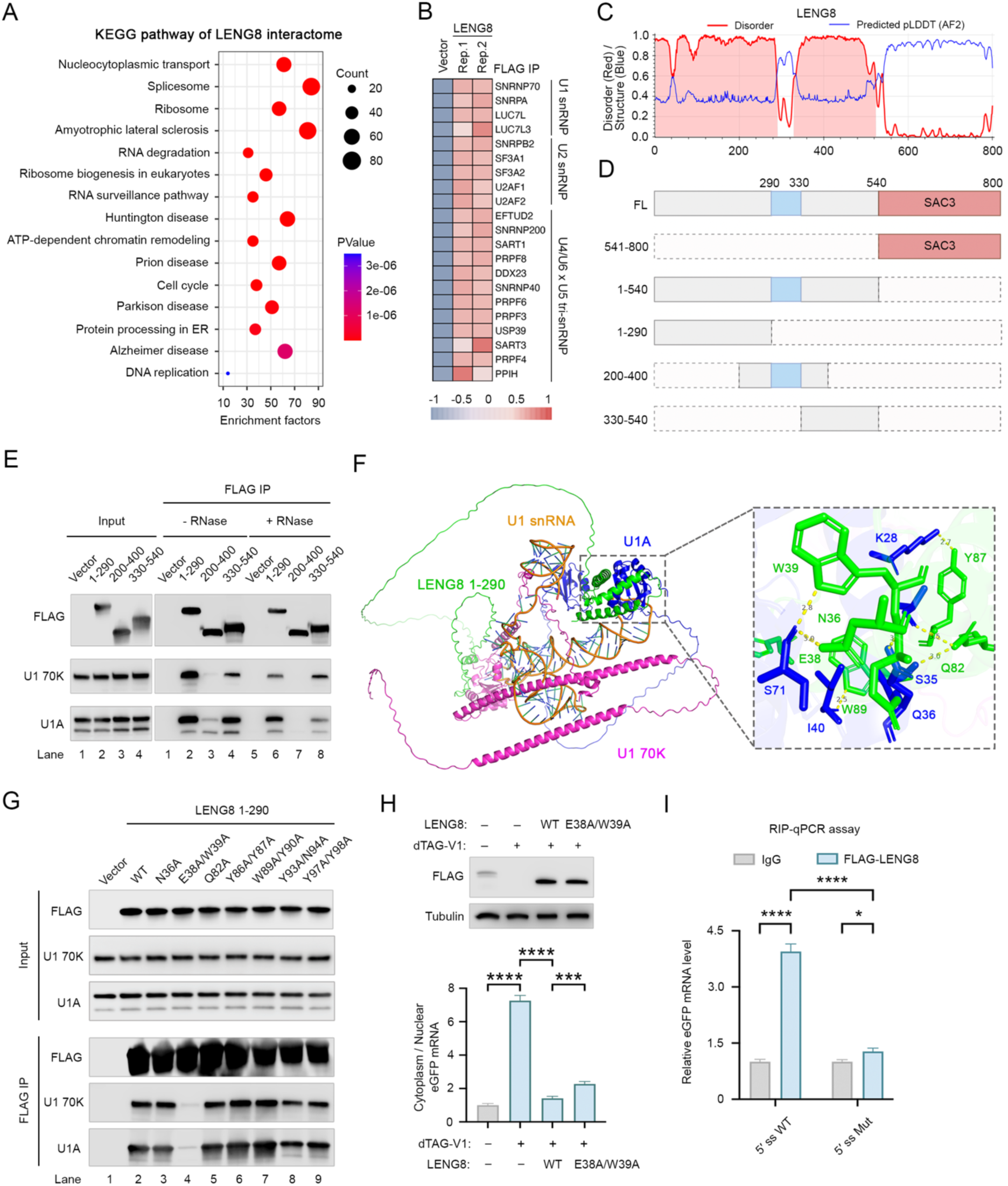
LENG8 is recruited to target RNAs via interactions with U1 snRNP and splicing factors. (A) KEGG pathway analysis of LENG8 interactors is performed using DAVID. The top 15 enriched pathways are displayed. (B) Heatmap showing LENG8-associated core spliceosome factors. (C) & (D) A schematic of LENG8 protein domains and the corresponding predicted disorder score based on ALBATROSS ^53^. (E) FLAG immunoprecipitation (IP) using different regions of LENG8, followed by western blot analysis. (F) Structure of the LENG8 (1-290aa)-U1 snRNP complex predicted by using AlphaFold3. The inset displays the specific amino acid residues involved in interactions. (G) FLAG immunoprecipitation (IP) using different LENG8 mutants (1-290aa), followed by western blot analysis. (H) A depletion and rescue assay. The top panel is western blot analysis to assess the rescue of LENG8 and LENG8 mutant (E38A/W39A) in degron-expressing cells treated with DMSO or dTAG-V1 for 24 hours. The bottom panel shows that the relative cytoplasmic eGFP mRNA level is measured in cells expressing either LENG8 or the LENG8 mutant (E38A/W39A) following LENG8 depletion by dTAG-V1, respectively. Cytoplasmic eGFP mRNA level is normalized to nuclear mRNA and quantified relative to the DMSO control condition. Data presented as mean ± SEM (n = 3). *p value % 0.05, **p value % 0.01, ***p value % 0.001, ****: p-value % 0.0001 (one-way ANOVA). (I) A RIP-PCR assay was performed to compare the 5′ ss WT or Mut RNA association with FLAG-LENG8. Data presented as mean ± SEM (n = 3). *p value % 0.05, **p value % 0.01, ***p value % 0.001, ****: p-value % 0.0001 (unpaired t-test).

To better understand the molecular basis of the interactions between LENG8 and U1 snRNP, we predicted the structure of a LENG8 (1-290aa)-U1 snRNP complex using AlphaFold3^39^. Interestingly, the predicted structure revealed two interactions between LENG8 and U1A: one mediated by 34-41aa and another by 81-102aa of LENG8 (Figure **4F**). And these two regions in LENG8 are highly conserved in mammals and mutating the conserved amino acid residues within these two regions significantly disrupted interaction between LENG8 (1-290aa) and the U1 snRNP (Figure **S4B, S4C**). The binding interfaces between LENG8 and U1 snRNP were quite extensive, involving many amino acid residues (Figure **4F**, inset). To disrupt the interactions between LENG8 and U1 snRNP specifically, we generated single or double mutations in LENG8 at the predicted binding interface, including N36A, E38A/W39A, Q82A, Y86A/Y87A, W89A/Y90A, Y93A/N94A, and Y97A/Y99A. We then expressed these mutant LENG8 proteins fused to a FLAG tag and performed FLAG IP. The results revealed that the E38A/W39A mutant displayed significantly weaker interactions with U1 snRNP (Figure **4G**, lane 4), suggesting that these two amino acid residues are required for interacting with U1 snRNP.

To test the functional significance of the LENG8-U1 snRNP interaction, we first depleted endogenous LENG8 protein in our degron cell line and then over-expressed the WT or a Mut LENG8 containing the E38A/W39A mutations. We then transfected the eGFP-5′ ss-PAS reporter into these cells and checked the reporter expression by measuring eGFP signals. Consistent with our earlier results, the reporter expression was low in control cells and LENG8 depletion led to a significant increase in eGFP signals. When the WT or Mut LENG8 was expressed in cells depleted of endogenous LENG8, the WT protein led to a more significant decrease in eGFP signal than the Mut (Figure **S4D**), suggesting that LENG8 interaction with U1 snRNP is required for its function in RNA nuclear retention and/or degradation. Furthermore, significantly higher cytoplasm/nucleus ratio was observed for the Mut LENG8 compared to WT protein, furthering supporting that LENG8 interaction with U1 snRNP is required for retaining 5′ ss-containing RNAs in the nucleus (Figure **4G**). Based on the LENG8-U1 snRNP interaction, we predicted that LENG8 is associated with 5′ ss-containing RNAs. Our RNA pulldown analyses already demonstrated that LENG8 associated with 5′ ss-containing RNA in vitro (Figure 1C). To test this in cells, we expressed the eGFP-5′ ss WT or Mut-PAS reporter and FLAG-LENG8, performed FLAG IP and examined the association between LENG8 and the reporter mRNA by RIP (RNA immunoprecipitation)-PCR analysis. Our results showed that LENG8 preferentially bound reporter mRNAs containing a 5′ ss (Figure **4I**, Figure **S4E**). Thus our results using in vitro and in-cell assays demonstrated that LENG8 is recruited to 5′ ss-containing RNAs via its interaction with the U1 snRNP and that this interaction is required for LENG8-mediated RNA nuclear retention and degradation.

### LENG8 represses RNA export by acting as a dominant negative factor of TREX-2

We next investigated the mechanism for LENG8-mediated nuclear retention of 5′ ss-containing RNAs. As mentioned earlier, the C-terminus of LENG8 contains a SAC3 domain, which is also found in the LENG8 homologue in yeast Thp3p (Figure 5A, Figure **S5A**). Previous studies have shown that Thp3 forms a stable complex with two other proteins, Csn12p and Sem1p^40^. Interestingly, our prediction using AlphaFold3 suggested that a similar complex is formed by the human homologues of these three proteins, LENG8, PCID2 and SEM1 and that LENG8 interactions with the other two proteins are mediated by its SAC3 domain (Figure **S5B, S5C, S5D, S5E**). Consistent with this, our LENG8 interactome analysis demonstrated a specific interaction between LENG8 and PCID2 (**Supplemental Table 4**). However, the function of this conserved LENG8-PCID2-SEM1 complex remains unknown.

**Figure 5.**
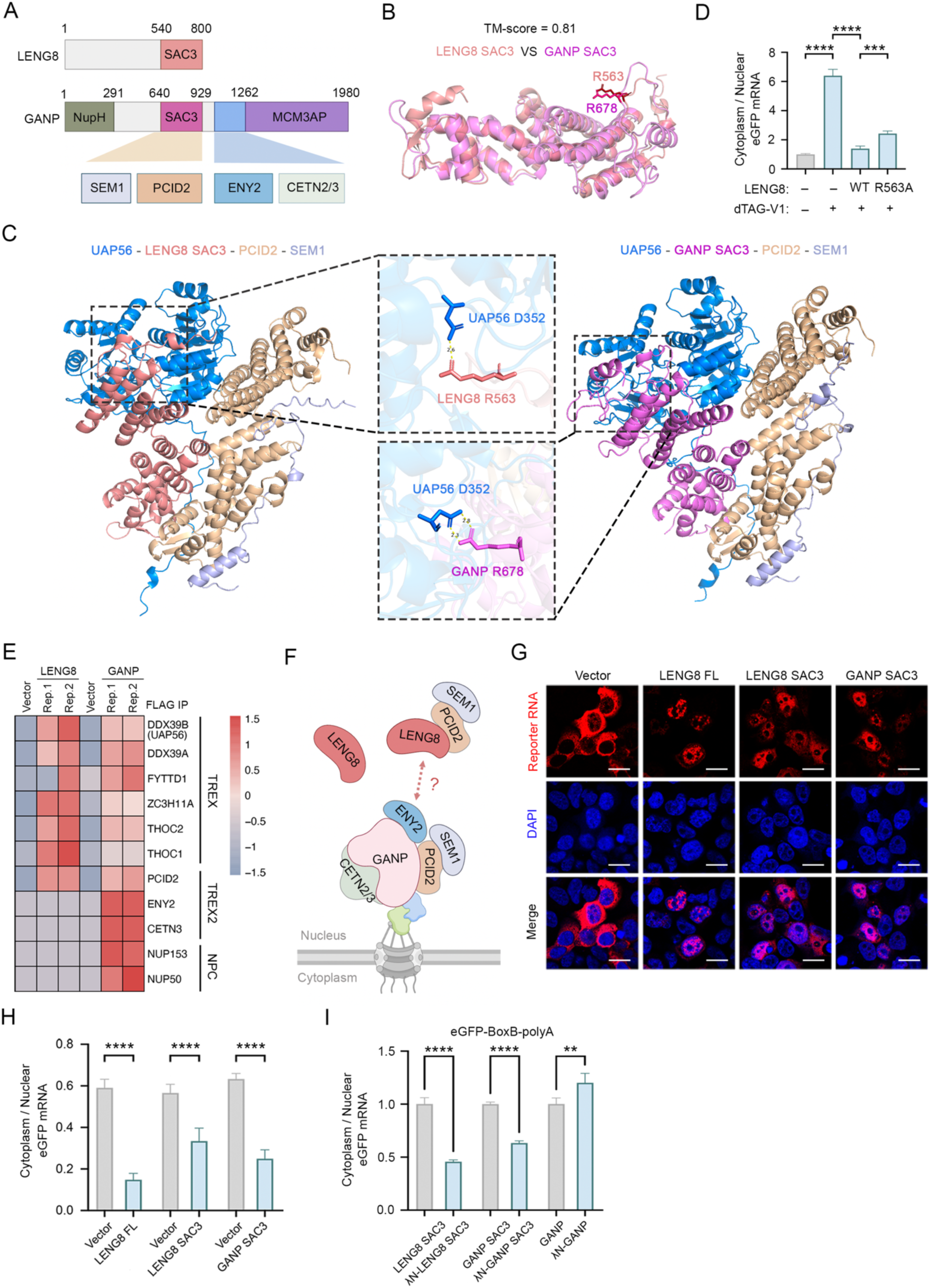
LENG8 represses RNA export by acting as a dominant negative factor of TREX-2. (A) Comparison of LENG8 and GANP protein domains. The domains and their interacting proteins are connected by colored boxes. (B) Comparison of LENG8 SAC3 and GANP SAC3 domains based on AlphaFold2 prediction and Pairwise Structure Alignment. The amino acid residue R678 in GANP SAC3 is equivalent to R563 in LENG8 SAC3. (C) Comparison of UAP56-LENG8 SAC3-PCID2-SEM1 and UAP56-GANP SAC3-PCID2-SEM1 complexes based on AlphaFold3 prediction. R678 in GANP SAC3 and R563 in LENG8 SAC3 both interact with UAP56 D352. (D) The relative cytoplasmic eGFP mRNA level is measured in cells expressing either LENG8 or the LENG8 mutant (R563A) following LENG8 depletion by dTAG-V1. Cytoplasmic eGFP mRNA level is normalized to nuclear mRNA and quantified relative to the DMSO control condition. Data presented as mean ± SEM (n = 3). *p value % 0.05, **p value % 0.01, ***p value % 0.001, ****: p-value % 0.0001 (one-way ANOVA).(E) Heatmap showing representative core factors of TREX, TREX-2, and NPC associated with LENG8 and GANP respectively. (F) A diagram for our model that LENG8 acts as a dominant-negative factor of GANP/TREX-2 as they compete for binding to PCID2 and SEM1. (G) RNA FISH analysis of eGFP reporter is performed after co-transfection of LENG8 FL, LENG8 SAC3 and GANP SAC3 construct, respectively. The eGFP reporter has no 5′ ss. The reporter RNA signal is shown in red, and DAPI is blue. Scale bars: 20 μm (H) The relative cytoplasmic eGFP mRNA level is measured in cells expressing eGFP reporter without 5′ ss after co-transfection of LENG8 FL, LENG8 SAC3 and GANP SAC3 construct, respectively. Cytoplasmic eGFP mRNA levels are normalized to nuclear mRNA and quantified relative to the negative control condition. Data presented as mean ± SEM (n = 3). *p value % 0.05, **p value % 0.01, ***p value % 0.001, ****: p-value % 0.0001 (unpaired t-test). (I) The relative cytoplasmic eGFP mRNA level is measured in cells co-expressing the eGFP-BoxB-polyA along with the SAC3 domain from LENG8 or GANP, either with or without an N-terminal λN peptide. Cytoplasmic eGFP mRNA level is normalized to nuclear mRNA and quantified relative to the no λN peptide control condition. Data presented as mean ± SEM (n = 3). *p value % 0.05, **p value % 0.01, ***p value % 0.001, ****: p-value % 0.0001 (unpaired t-test). (J) A RIP-PCR assay is performed to compare the RNA binding preferences of GANP. Data presented as mean ± SEM (n = 3). *p value % 0.05, **p value % 0.01, ***p value % 0.001, ****: p-value % 0.0001 (unpaired t-test).

Interestingly, the SAC3 domain is also found in GANP, a core component of the essential RNA export factor TREX-2^41,42^ (Figure **5A**). The structures of the SAC3 domains of LENG8 and GANP predicted by using AlphaFold3 showed high degree of similarity (TM-score = 0.83) (Figure **5B**). Similar to LENG8, GANP also forms a stable complex with PCID2 and SEM1 and the two trimeric complexes are again highly similar based on our prediction (Figure **5C**) and the recently published cryo-EM structures^43,44^. In addition, a recent study demonstrated that the GANP/PCID2/SEM1 complex can bind to UAP56 and stimulates its ATPase activity, thereby facilitating its release from the mRNA, and that this activity is required for mRNA export^44,45^. This interaction is mediated by the R678 of GANP and D352 of UAP56, and mutating R678 in GANP abolished its activation of UAP56 ATPase activity and caused mRNA export defect. Interestingly AlphaFold3 predicted a similar binding pattern between the LENG8/PCID2/SEM1 complex and UAP56 (Figure **5C**, left panels). Specifically, R563 in LENG8, which is conserved from yeast to human, interacts with D352 of UAP56 (Figure **5C**, middle panels, Figure **S6A, S6B, S6C**). To test the functional significance of this interaction, we depleted endogenous LENG8 and rescued with either the WT or a R563A mutant LENG8 and checked its impact on the eGFP-5′ ss-PAS reporter mRNA export. Our results showed that the R563A mutant was significantly less active compared to the WT in retaining the reporter mRNAs in the nucleus, suggesting that the LENG8-UAP56 interaction is important for LENG8 function in nuclear retention (Figure **5D**, Figure **S6D**).

There are also major differences between LENG8 and GANP. Although they both contain SAC3 domain, LENG8 lacks the other domains in GANP that mediate critical interactions with other TREX-2 and NPC components that are required for mRNA export (Figure **5A**). Indeed, when we compared the interactomes of LENG8 and GANP through FLAG IP and MS analyses (**Supplemental Table 4, 6**), we found that both proteins bind to TREX and PCID2, but LENG8 failed to interact with TREX-2 components such as ENY2 and CETN3, or NPC components NUP153 and NUP50 (Figure **5E**), both of which are essential for anchoring GANP and TREX-2 to the NPC^46^. These data suggests that LENG8 may function as a dominant negative factor for TREX-2 by competing with GANP for binding to TREX, UAP56, and PCID2/SEM1 (Figure **5F**). This model would predict that over-expressing LENG8 or its SAC3 domain itself could repress mRNA export. To test this, we over-expressed the FL or the SAC3 domain of LENG8 or GANP and monitored their effect on the mRNA localization by RNA FISH for a reporter that contained eGFP coding sequence followed by PAS (Figure **5G**). In control cells expressing an empty vector, the reporter mRNAs were predominantly localized to the cytoplasm. In contrast, over-expression of FL LENG8 or the SAC3 domains of LENG8 or GANP led to nuclear retention of the reporter mRNAs (Figure **5G**, quantification in Figure **5H**). In addition to this reporter, we also monitored impact of over-expressing LENG8 or SAC3 domains on the localization of cellular bulk mRNAs by performing RNA FISH using an oligo(dT) probe and observed a similar pattern (Figure **S6E**), providing evidence that indeed LENG8 functions as a dominant negative factor for TREX-2.

Our earlier data demonstrated that tethering the FL LENG8 to reporter mRNAs is sufficient to retain them in the nucleus (Figure **2F, 2H**). We next tested the effect of tethering the SAC3 domains of LENG8 or GANP to reporter mRNAs on their localization. Our results showed that tethering the SAC3 domain was sufficient to inhibit reporter mRNA export, while tethering FL GANP had the opposite effect (Figure **5I**, Figure **S6F**), providing further evidence that, similar to the SAC3 domain alone, LENG8 can inhibit mRNA export by acting as a dominant negative factor to GANP/TREX-2. According to our model, LENG8 functions as a dominant negative factor of GANP/TREX-2 because it interacts with PCID2/SEM1, but cannot interact with ENY2/CENT3 and other GANP-associated factors. This model would predict that fusing the GANP domain mediating interactions with ENY2/CENT3 to LENG8 SAC3 domain would alleviate its inhibitory activity on mRNA export. To test this, we constructed a chimera by fusing GANP 990-1350aa with the SAC3 domain of LENG8 (Figure **S6G**). Structural prediction by Alphafold3 suggested that this chimera can recruit ENY2 and CENT3 and we verified this interaction by FLAG IP (Figure **S6G, S6H**). When this chimera was tethered to the reporter, we found that it had significantly lower inhibitory activity on the reporter expression than LENG8 SAC3 domain alone (Figure **S6I**). Together, these findings suggest that LENG8 retains misprocessed RNAs in the nucleus by interfering with GANP/TREX-2 function in a dominant-negative manner.

### LENG8 facilitates RNA degradation via interactions with the PAXT and exosome complexes

Several lines of evidence suggest that LENG8 mediates RNA degradation by recruiting the RNA exosome adaptor PAXT and the RNA exosome. First, PAXT components, ZFC3H1 and MTREX/MTR4, and the exosome subunits were among the top hits from our CRISPR screen (Figure **1**). Secondly, our interactome analysis showed that LENG8 associated with PAXT and the exosome (Figure **6A**). Thirdly, previous studies, including ours, showed that PAXT recognizes IPA transcripts and PROMPTs for exosome-mediated degradation^14,16^. In this study, we showed that LENG8 also specifically targets IPA transcripts and PROMPTs for degradation (Figure **2** **and** Figure **3**). Importantly, when we compared the IPA transcripts targeted by PAXT, LENG8, and the exosome by hierarchical clustering, we found that LENG8 clustered closely with PAXT components ZFC3H1 and MTR4 (Figure **6B**). As shown in Figure **6C**, the IPA transcripts of the *TP53* and *PCF11* gene significantly increased upon depletion of LENG8, ZFC3H1, MTR4, and EXOSC3, further supporting that LENG8 and PAXT mediate RNA degradation in the same pathway.

**Figure 6.**
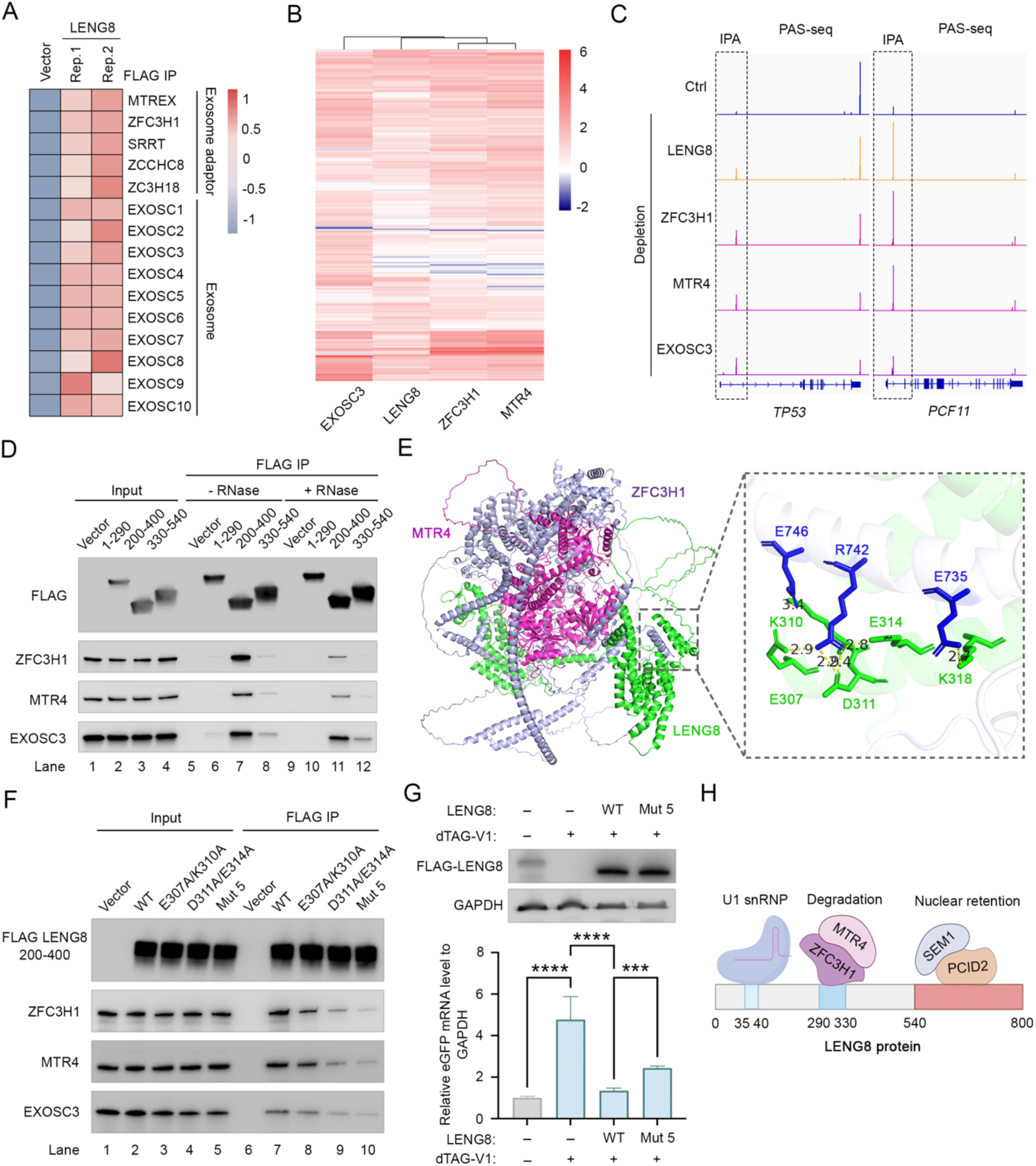
LENG8 facilitates RNA degradation via interactions with PAXT and the RNA exosome. (A) Heatmap showing representative LENG8-associated core RNA exosome adaptors and all of exosome factors. (B) Heatmap of the log2FC measured for all IPA transcripts that are upregulated after depletion of EXOSC3, LENG8, ZFC3H1, and MTR4 (log2FC > 1, FDR ≤ 0.05). (C) PAS-seq tracks showing the *TP53* and *PCF11* following depletion of LENG8, ZFC3H1, MTR4 and EXOSC3. IPA signals are marked by boxes. (D) FLAG immunoprecipitation (IP) using different regions of LENG8, followed by western blot analysis. (E) The structure of LENG8(200-400aa)- ZFC3H1-MTR4 complex predicted by using AlphaFold3. (F) FLAG immunoprecipitation (IP) using different LENG8 (200-400aa) mutants, followed by western blot analysis. (G) LENG8 depletion and rescue assay. The top panel is western blot analysis to assess the rescue of LENG8 and LENG8 mutant (E307A/K310A/D311A/E314A/KK318A) in degron-expressing cells treated with DMSO or dTAG-V1 for 24 hours. The bottom panel showed that eGFP mRNA level is measured in cells expressing either LENG8 or the LENG8 mutant (E307A/K310A/D311A/E314A) following LENG8 depletion by dTAG-V1. The relative eGFP mRNA level is normalized to co-transfected mCherry and then quantified relative to the DMSO control condition. Data presented as mean ± SEM (n = 3). *p value % 0.05, **p value % 0.01, ***p value % 0.001, ****: p-value % 0.0001 (one-way ANOVA). (H) A schematic summarizing interactions between LENG8 and its key partners.

We next characterized the interactions between LENG8 and PAXT components, ZFC3H1 and MTR4, by performing FLAG IP with the FL or truncated LENG8. Our results showed that the FL and the N-terminal disordered region, but not the C-terminal SAC3/GANP/THP3 domain, of LENG8 interacted with PAXT (Figure **S7A**). When examining different fragments within the N-terminal region, we found that the fragment consisting of 200-400aa specifically interacted with ZFC3H1-MTR4-exosome complex, but not the other fragments (Figure **6D**, lanes 7 and 11). This region is distinct from the region required for binding to U1 snRNP (1-290aa and 330-540aa, Figure **4E**). In addition, we found that 290-330aa is also highly conserved in mammals and that alanine substitution of 290-330aa abolished the interaction between LENG8 (200-400aa) and PAXT (Figure **S7B, S7C**). To examine the interactions between LENG8 and PAXT in greater detail, we predicted the structure of the LENG8(200-400aa)-PAXT complex (Figure **6E**). The structure revealed that residues 290-330 of LENG8 form an α-helix and mediate interactions with ZFC3H1 via a few key residues, including E307, K310, D311, E314, and K318 (Figure **6E**). To test these predictions, some or all of these residues were mutated to alanine and the FLAG-tagged WT or mutant LENG8 fragments were expressed in HEK293T cells and FLAG IP assays were performed. The results showed that both the E307A/K310A and D311A/E314A had diminished binding to ZFC3H1 and MTR4 while mutating all 5 residues had the most significant effect (Figure **6F**). Reciprocally, mutations E735A/R742A/E746A in ZFC3H1 also decreased its interaction with LENG8 (Figure **S7D**), confirming the predicted interactions. To assess the function of LENG8-PAXT interaction in LENG8-mediated RNA degradation, a depletion and rescue assay was performed. As expected, overexpression of wild-type LENG8 in LENG8-depleted cells rescued the RNA degradation of the eGFP-5′ ss WT-PAS reporter (Figure **6G**). In contrast, cells expressing the mutant LENG8 (Mut5) deficient in PAXT-binding exhibited significantly weaker activity compared to WT LENG8 (Figure **6G**), suggesting that LENG8 interaction with ZFC3H1-MTR4-exosome is required for efficient RNA degradation. Together, our results revealed that LENG8 is a modular protein with distinct fragments mediating interaction with U1 snRNP, PCID2/SEM1, and PAXT (Figure **6H**). Through these interactions, LENG8 is specifically recruited to 5’ ss-containing RNAs, including IPAs and intron-retained mRNAs, and mediate their nuclear retention and degradation.

## Discussion

It has long been known that incompletely processed or misprocessed RNAs are retained in the nucleus^10,11^, but the mechanism remains unknown. Although several factors have been implicated in this process, including splicing factors (U1 snRNP)^15,25,47^, structural components of the NPC (Mpl1p/TRP) or degradation factors (ZFC3H1)^15,18,48^, it is unclear whether their putative function in nuclear retention is direct or indirect. In this study, we identified LENG8 as a key nuclear retention factor (Figure 7). We show that LENG8 is recruited to pre-mRNAs via splicing factors, including U1 snRNP, and is both necessary and sufficient to sequester bound RNAs in the nucleus. Mechanistically, LENG8 forms a complex with PCID2 and SEM1, and blocks mRNA export by acting as a dominant negative factor for TREX-2. Upon splicing completion and the dissociation of the splicing machinery, LENG8 also dissociates from RNAs, allowing the transfer of mRNAs to TREX-2 complex for export. For misprocessed RNAs with unused 5′ ss (e.g. intron-retained RNAs or IPAs), LENG8 mediates their nuclear retention. If the 5′ ss is followed by a poly(A) junction (e.g. in IPA transcripts), LENG8 promotes the recruitment of PAXT and the RNA exosome for degradation (Figure 7). Thus LENG8 plays a central role in determining the fate of RNAs based on their processing status.

**Figure 7.**
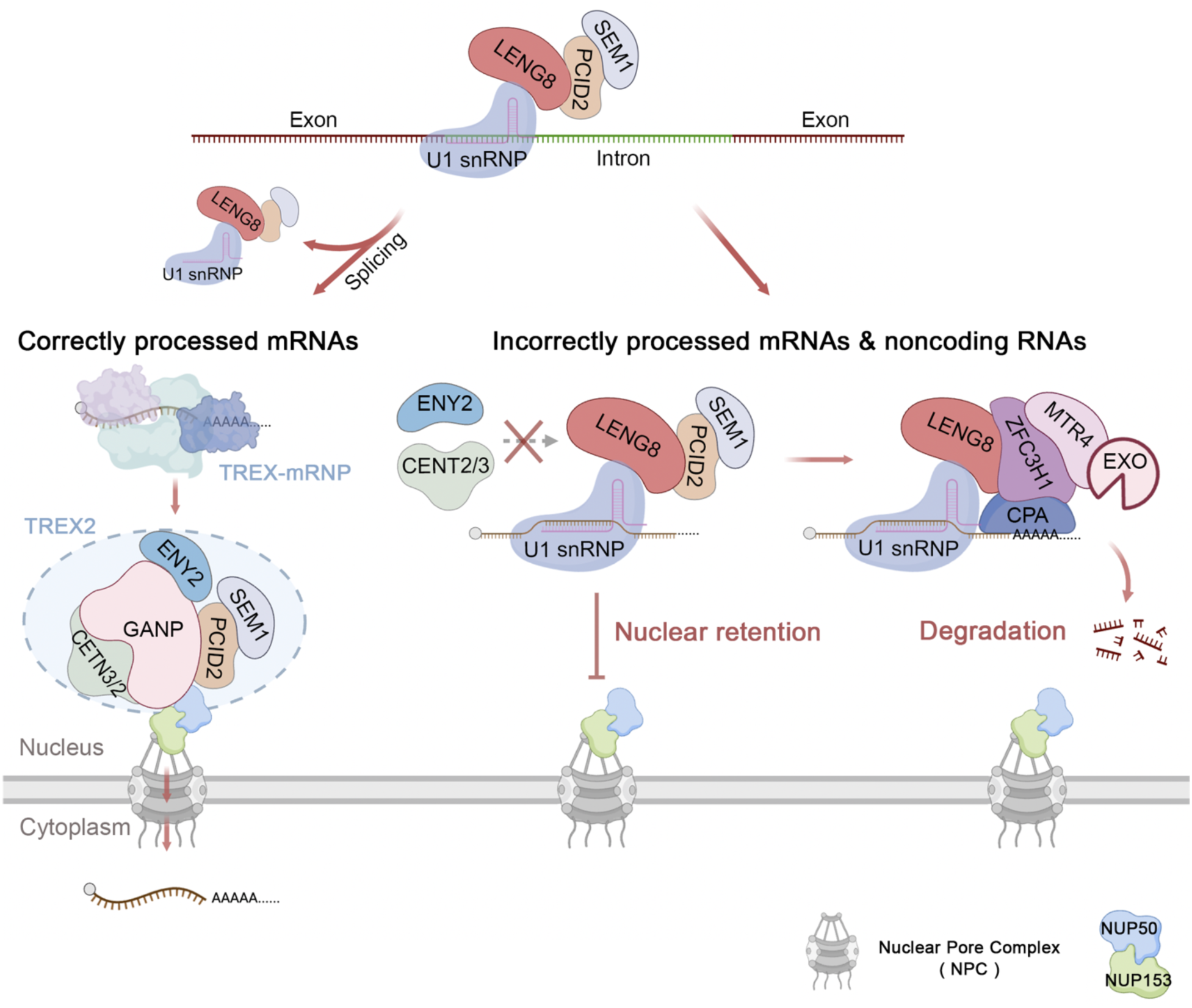
A model for LENG8-mediated RNA nuclear retention and degradation. A schematic model depicting the function of LENG8 as a key quality control factor that determines the fate of RNAs based on their processing status. Please see Discussion for details

For over 30 years, 5′ ss and the U1 snRNP are known to cause RNA nuclear retention from yeast to human^23–25^, but the mechanism has remained elusive. Two models have been proposed for nuclear retention: inefficient recruitment of export factors or recruitment of nuclear retention factors^10,20^. In this study, we provided strong evidence for the second model by identifying LENG8 as a 5′ ss-dependent and U1 snRNP-associated nuclear retention factor. LENG8 forms a complex with PCID2 and SEM1 and it blocks RNA export by acting as a dominant negative factor for TREX-2. Given the function of this complex in repressing RNA export, we refer to it as the REX (Repressor of Export) complex. Importantly all three subunits of the REX complex are conserved from yeast to human and the yeast complex has been linked to RNA surveillance and export^40^, suggesting that REX may be a conserved nuclear retention factor. Interestingly, we showed that LENG8 can sequester IPA or intron-retained RNAs in the nucleus even when multiple other introns in the same transcript have already been spliced, and thus EJCs have been deposited (Figure 3C, 3F). This observation suggests that the nuclear retention activity of LENG8 dominates the activities of the export factors. Although the underlying mechanism is currently unknown, one possibility is that LENG8 itself is anchored to certain nuclear structures, rendering it immobile. Thus, one RNA-bound LENG8 is sufficient to retain this RNA in the nucleus. Alternatively, our data demonstrated that both REX and TREX-2 complexes bind to UAP56 and stimulate UAP56 removal from mRNAs. If REX removes multiple or all UAP56 from an RNA molecule, this will block the recruitment of TREX-2, rending the RNA incompetent for export. It will be critical to test these models to fully understand how LENG8 mediates RNA nuclear retention.

Both IPAs and intron-retained mRNAs are sequestered in the nucleus in a LENG8-dependent manner, but the former group are also degraded while the stability of the latter group does not seem be influenced by LENG8. How are their different fate determined? Although LENG8 mediates 5′ ss-dependent recruitment of PAXT and exosome, our recent study demonstrated that this interaction is weak and requires CPA factors bound at the poly(A) junction to synergistically recruit PAXT/exosome for degradation. This is exactly what happens to IPA transcripts. On the other hand, as intron-retained RNAs contain both 5′ and 3′ ss, thus the RNA-bound U1 snRNP is likely engaged with U2AF and U2 snRNP, which prevents synergistic interactions between U1 snRNP and the CPA machinery. Thus intron-retained RNAs are retained in the nucleus, but not degraded. Intriguingly, upon LENG8 depletion, intron-retained RNAs that are leaked into the cytoplasm are highly enriched in the last introns (Figure 3F and Figure S3E). This could be due to the fact that the inclusion of upstream introns is more likely to induce nonsense-mediated RNA decay^49^. Alternatively, nuclear retention of RNAs containing unspliced internal introns may require a distinct and LENG8-independent mechanism.

In our study, we have demonstrated that LENG8 can be recruited to 5′ ss-containing RNAs via interactions with the U1 snRNP (Figure **4E-I**). Additionally, LENG8 also binds to other splicing factors, including U2AF1/2, U2 snRNP, and the U4/5/6 tri-snRNP (Figure **4B** and Figure **S4A**), raising the possibility that LENG8 could bind to RNA through other splicing factors as well. Indeed, previous studies have reported that U2AF65 binding could lead to RNA nuclear retention^47^, and it is possible that U2AF65-mediated RNA nuclear retention also requires LENG8. Furthermore, our data showed dramatic PROMPT RNAs accumulation and export into the cytoplasm in LENG8-depleted cells (Figure **3G-I**). Previous studies have suggested that 5′ ss are depleted in PROMPTs and these RNAs are generally not spliced^50^. Thus, LENG8 recruitment to these RNAs is likely independent of U1 snRNP or other splicing factors. Interestingly, our study detected interactions between LENG8 and the Restrictor complex, consisting of WDR82 and ZC3H4 (**Supplemental Table 4**), which has recently been shown to play an important role in regulating the biogenesis and degradation of PROMPTs and other noncoding RNAs^51,52^. It will be interesting to determine the functional links between LENG8 and the Restrictor complex in future studies.

## Data Availability

mRNA-seq and PAS-seq datasets have been deposited into the GEO under accession number GSE299756. Proteome datasets have been deposited into the ProteomeXchange under accession number PXD065356.

## Author Contributions

Conceptualization: L.T. and Y.S.; Investigation: L.T. and Y.S.; Formal Analysis: L.T. and Y.S.; Methodology: L.T., L.L., Y.Y., L.V.S., C.Y.; Original Draft: L.T. and Y.S.; Review & Editing: L.T. and Y.S.; Funding Acquisition: L.T. and Y.S.; Supervision: L.H. and Y.S.

## Acknowledgments

We wish to acknowledge the support of the Chao Family Comprehensive Cancer Center Shared Resource Genomics High-Throughput Facility, and the support of the National Cancer Institute of the National Institutes of Health under award number P30CA062203. We would like to thank Seung-Ah Yoon from the UCI Genomics High-Throughput Facility, and Pauline Nguyen from the UCI Stem Cell Center for technical support. This study was supported by the following grants: R35GM149294 (Y.S.) and R35GM145249 (L.H.). L.T. was supported by the Hewitt Foundation Postdoctoral Fellowship.

## STAR★Methods

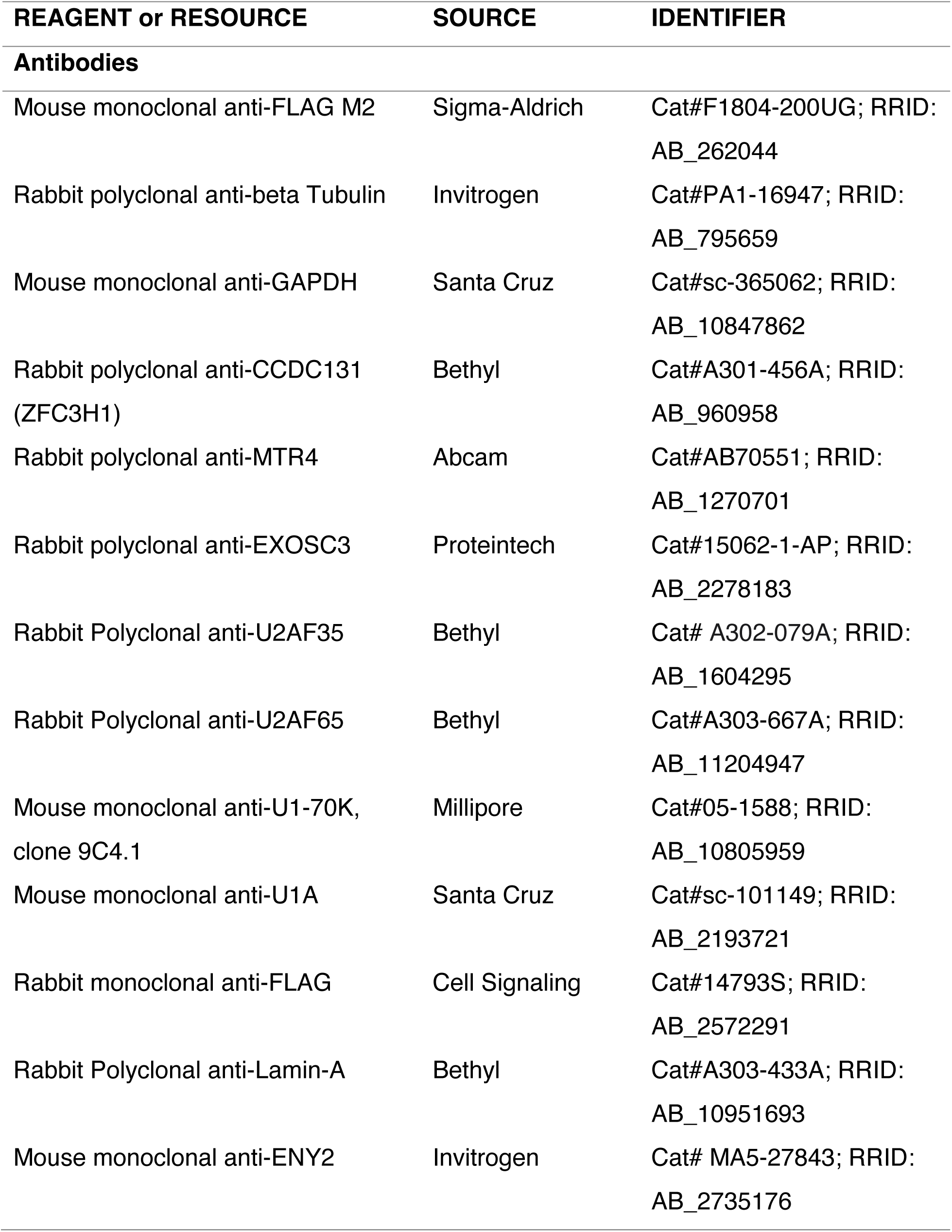
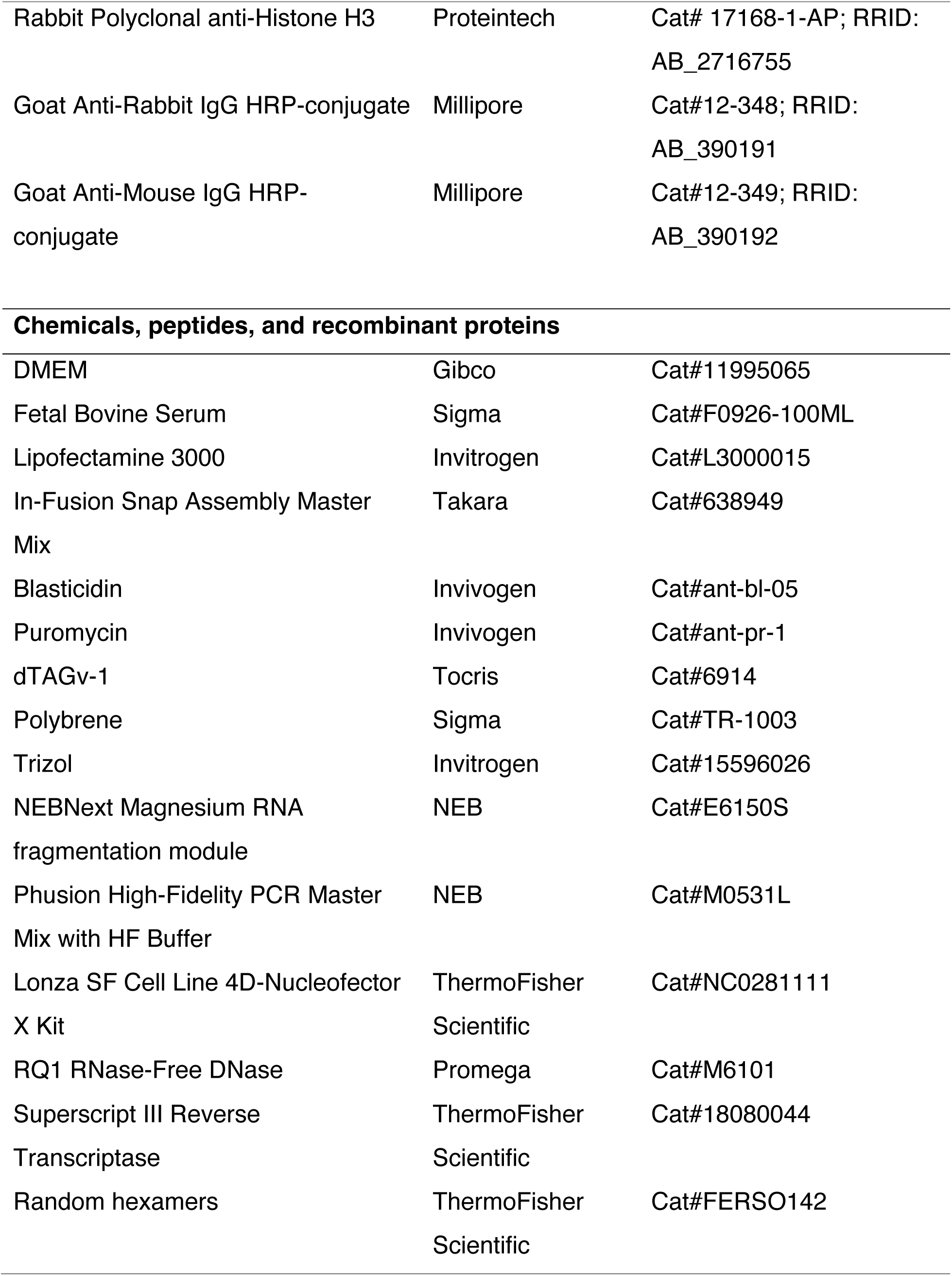
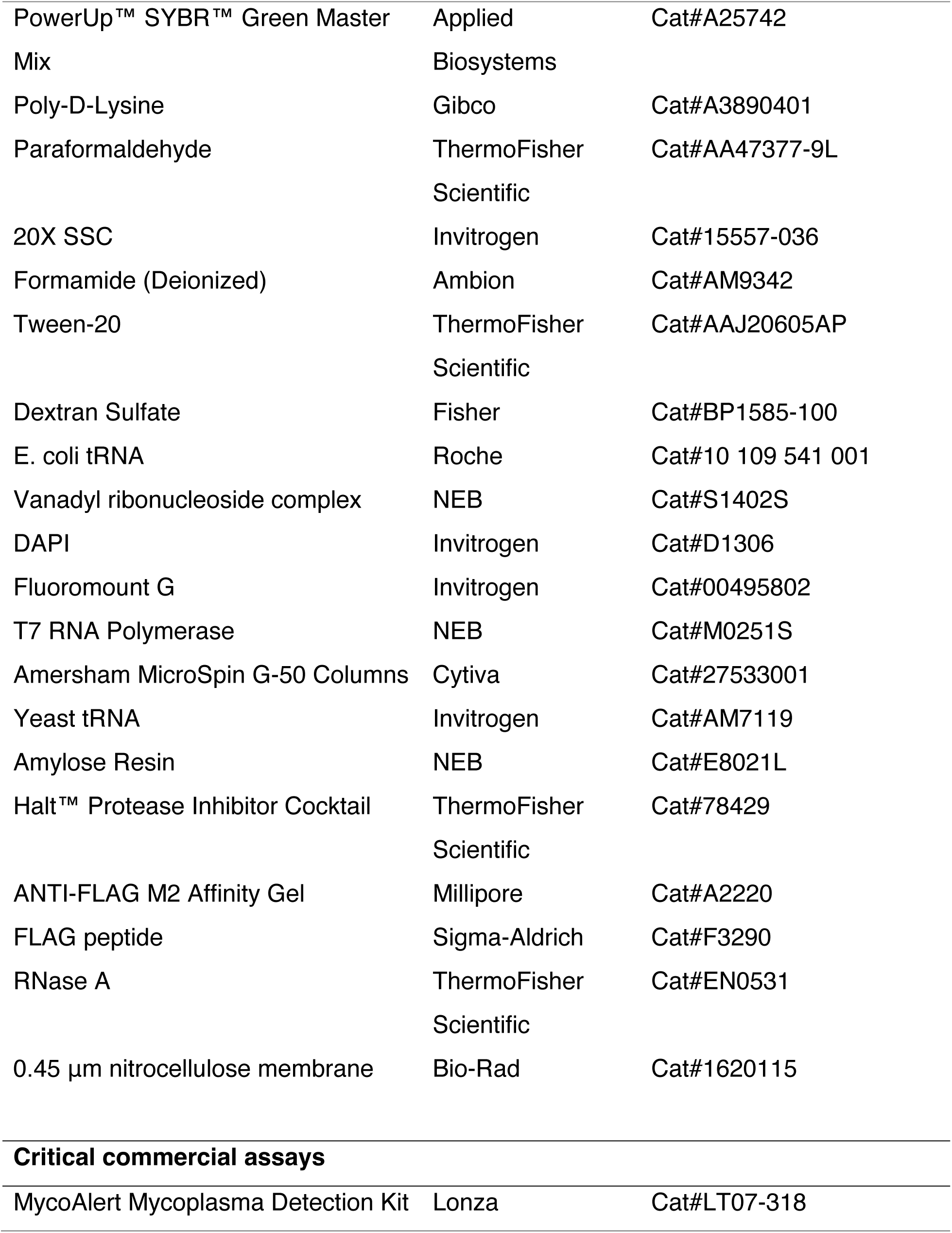
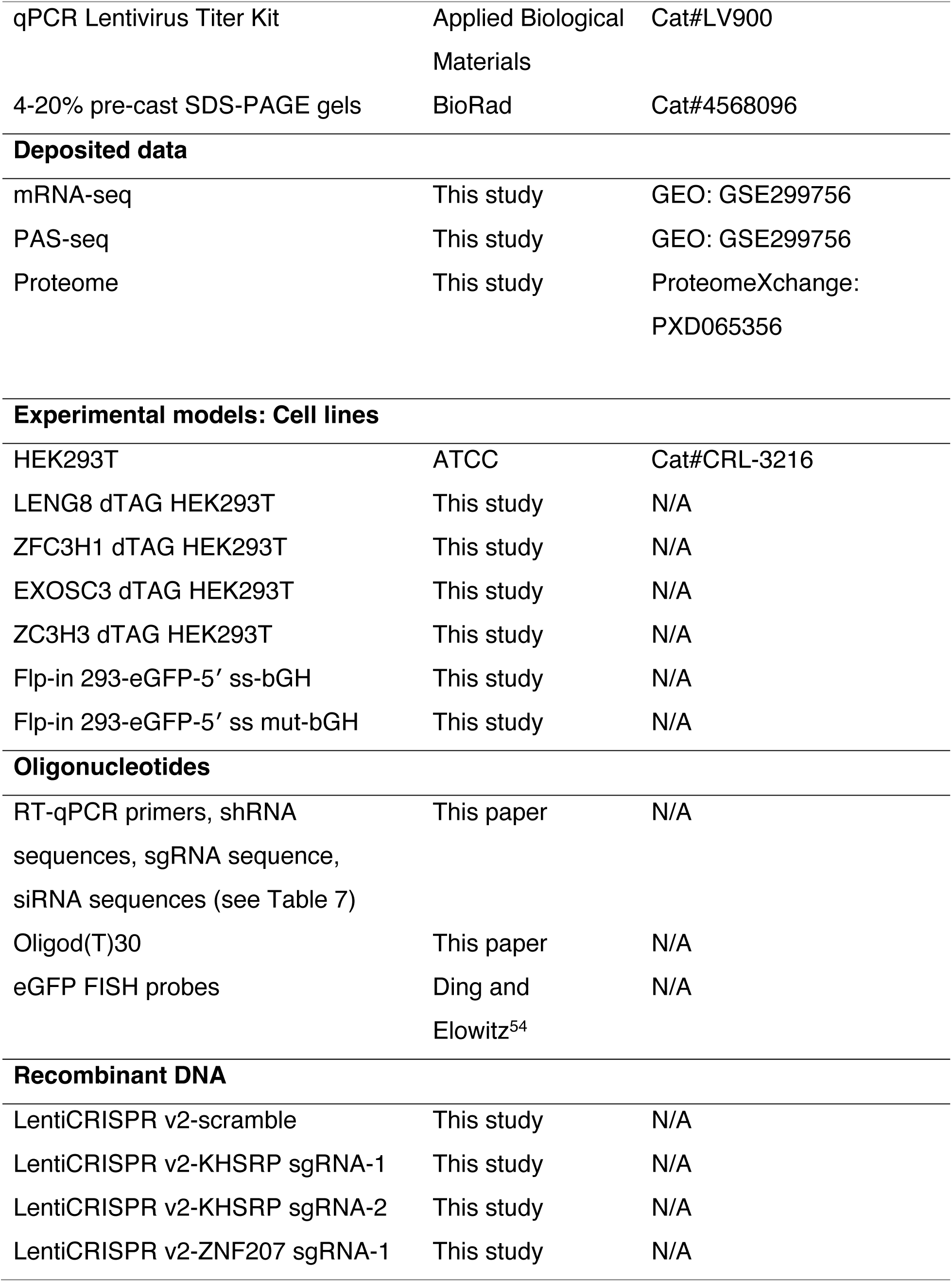
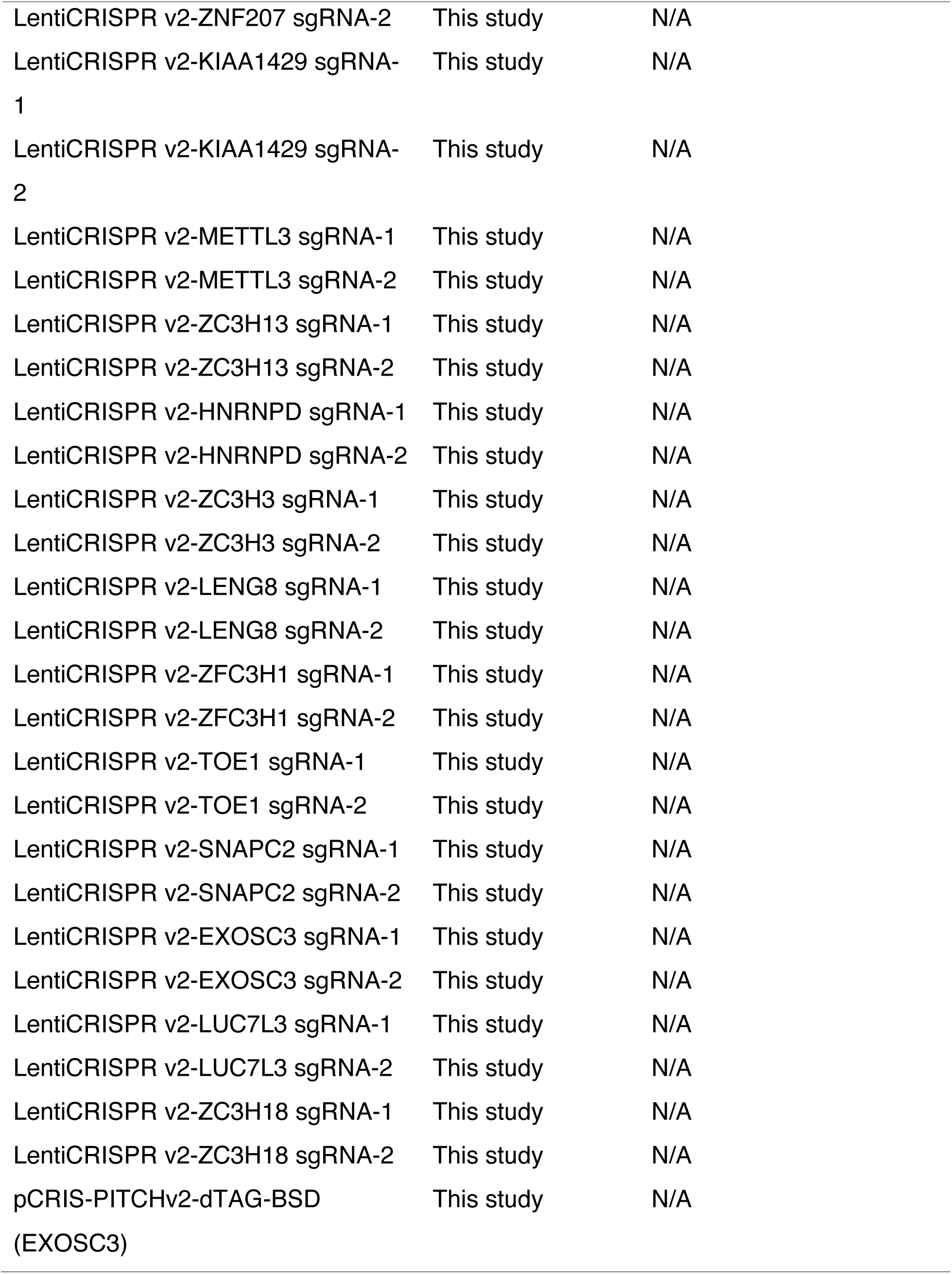
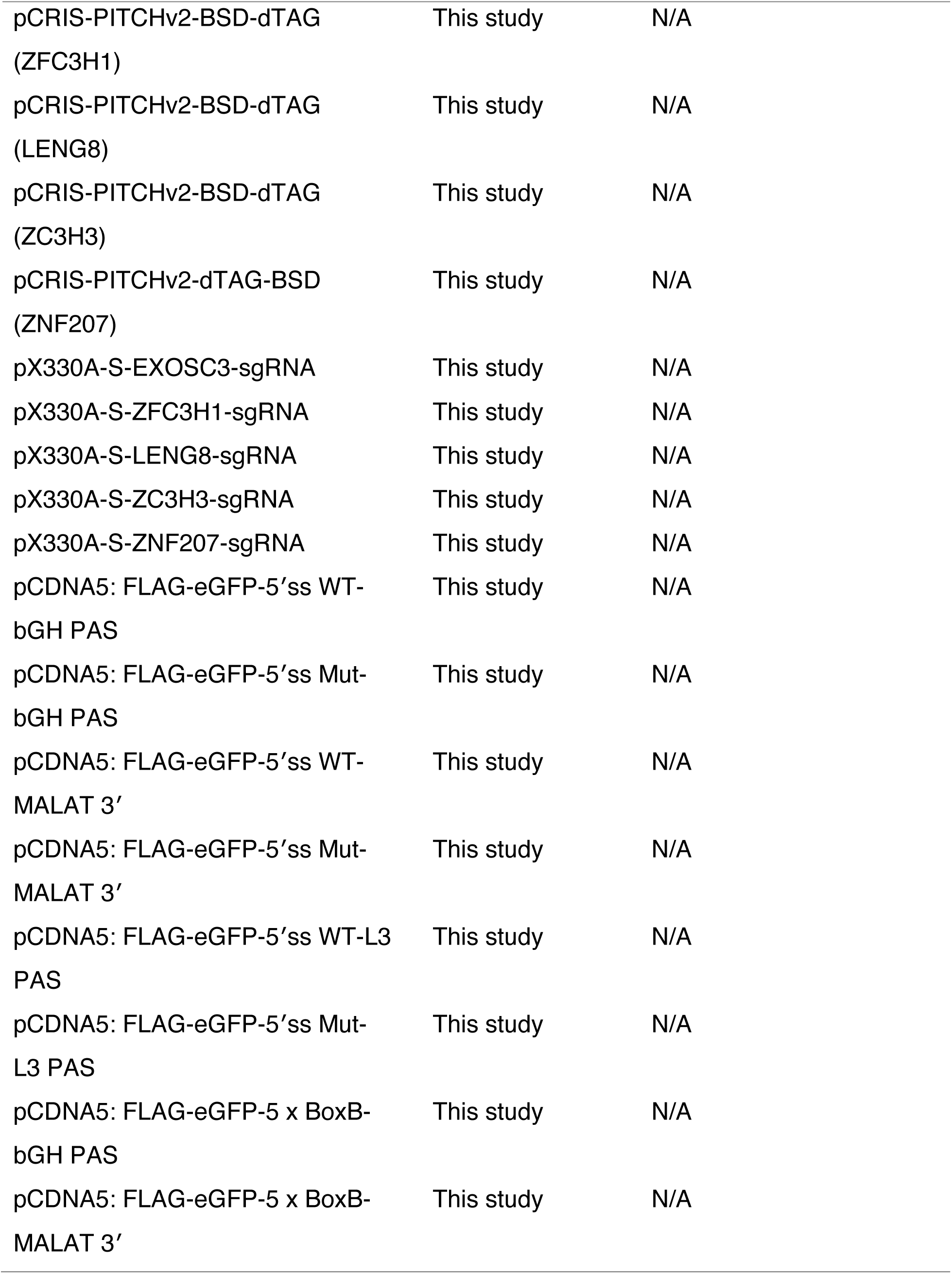
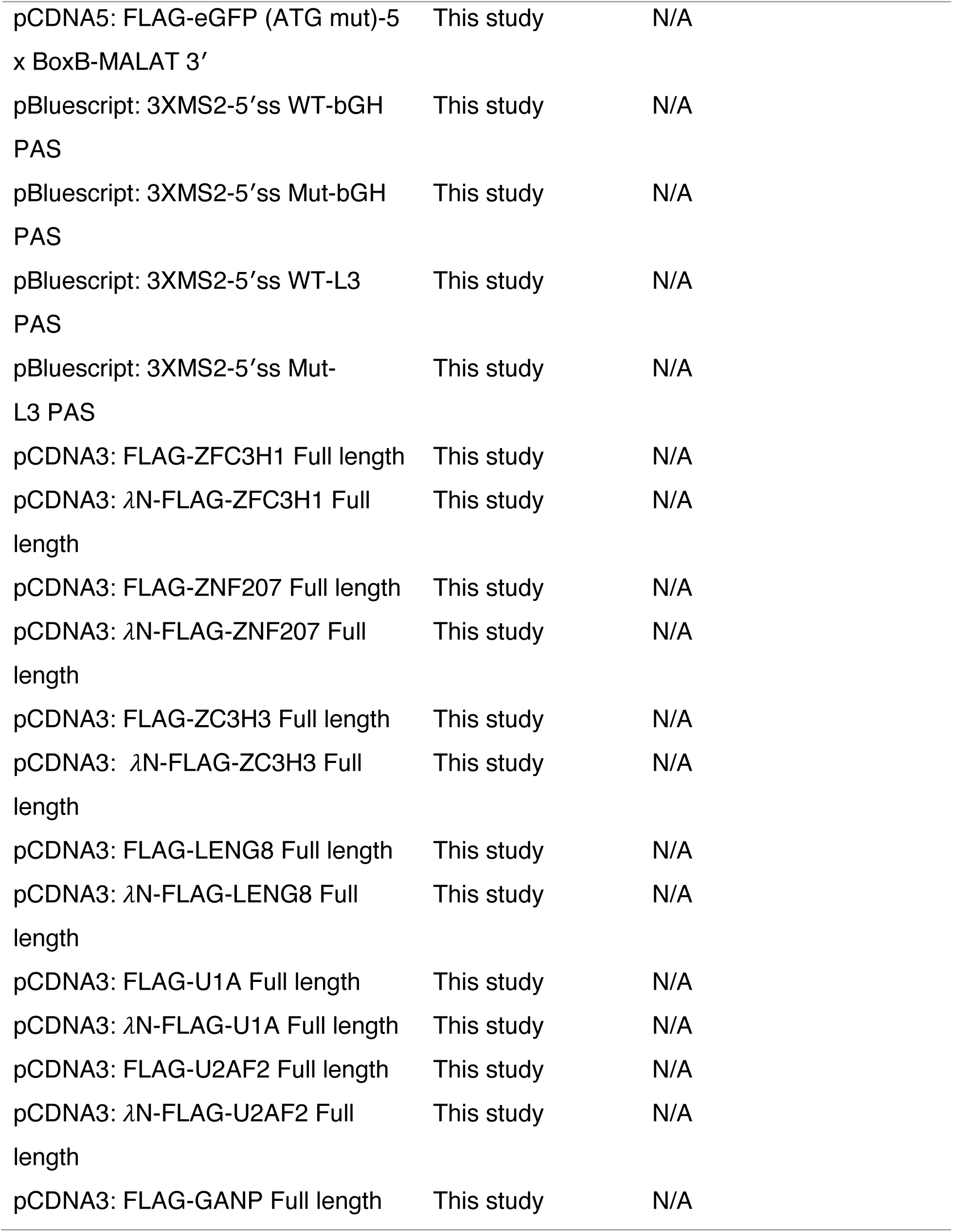
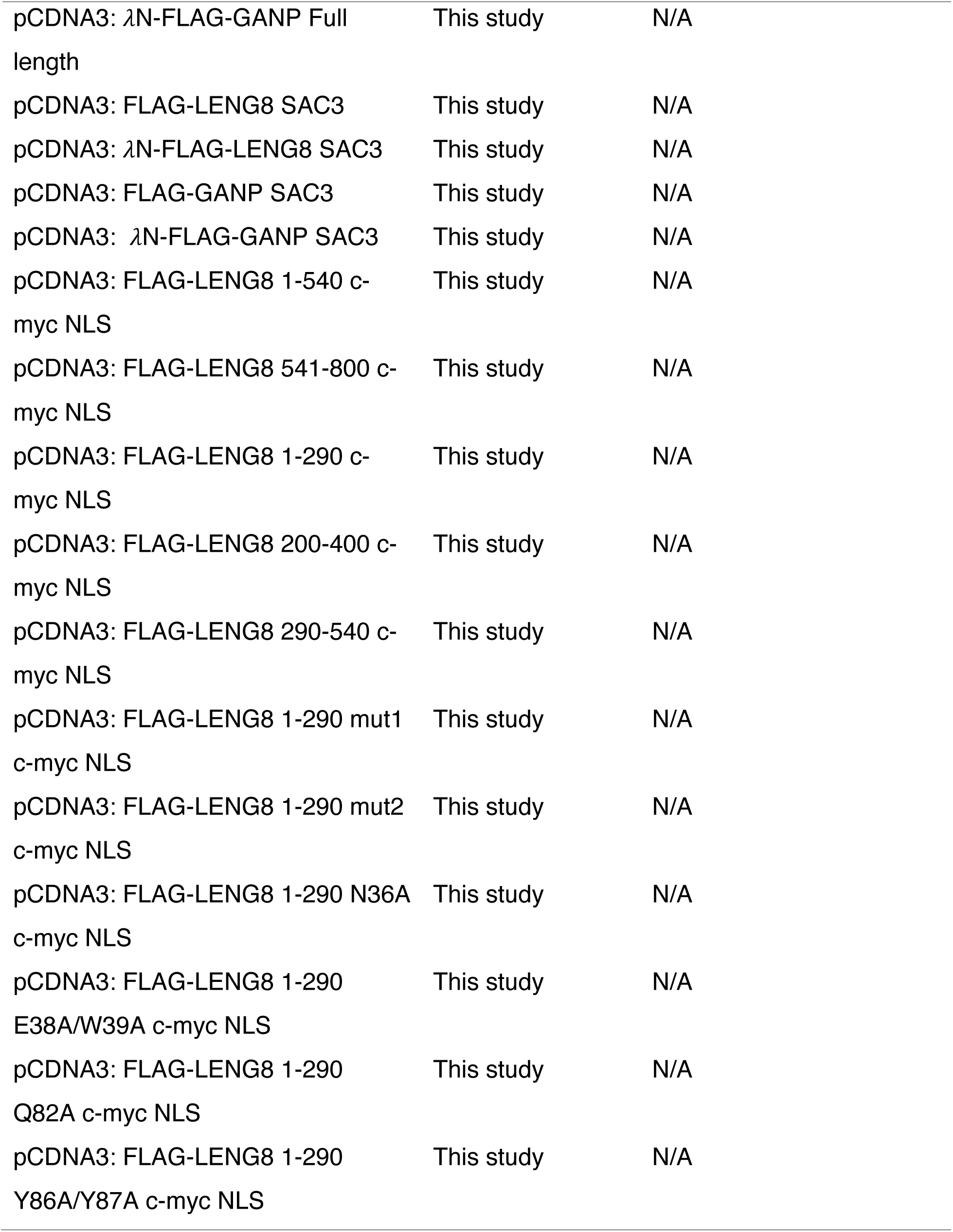
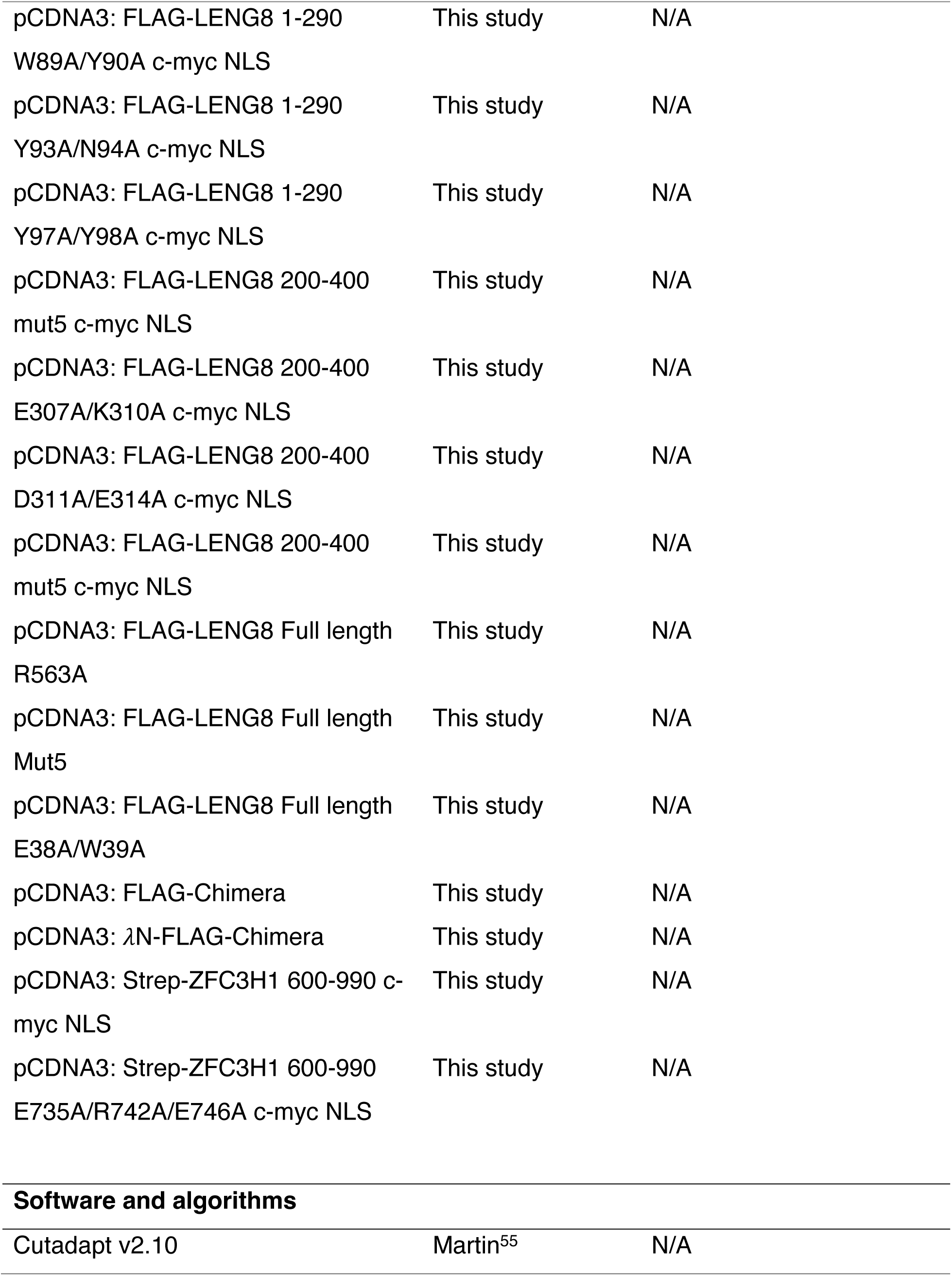
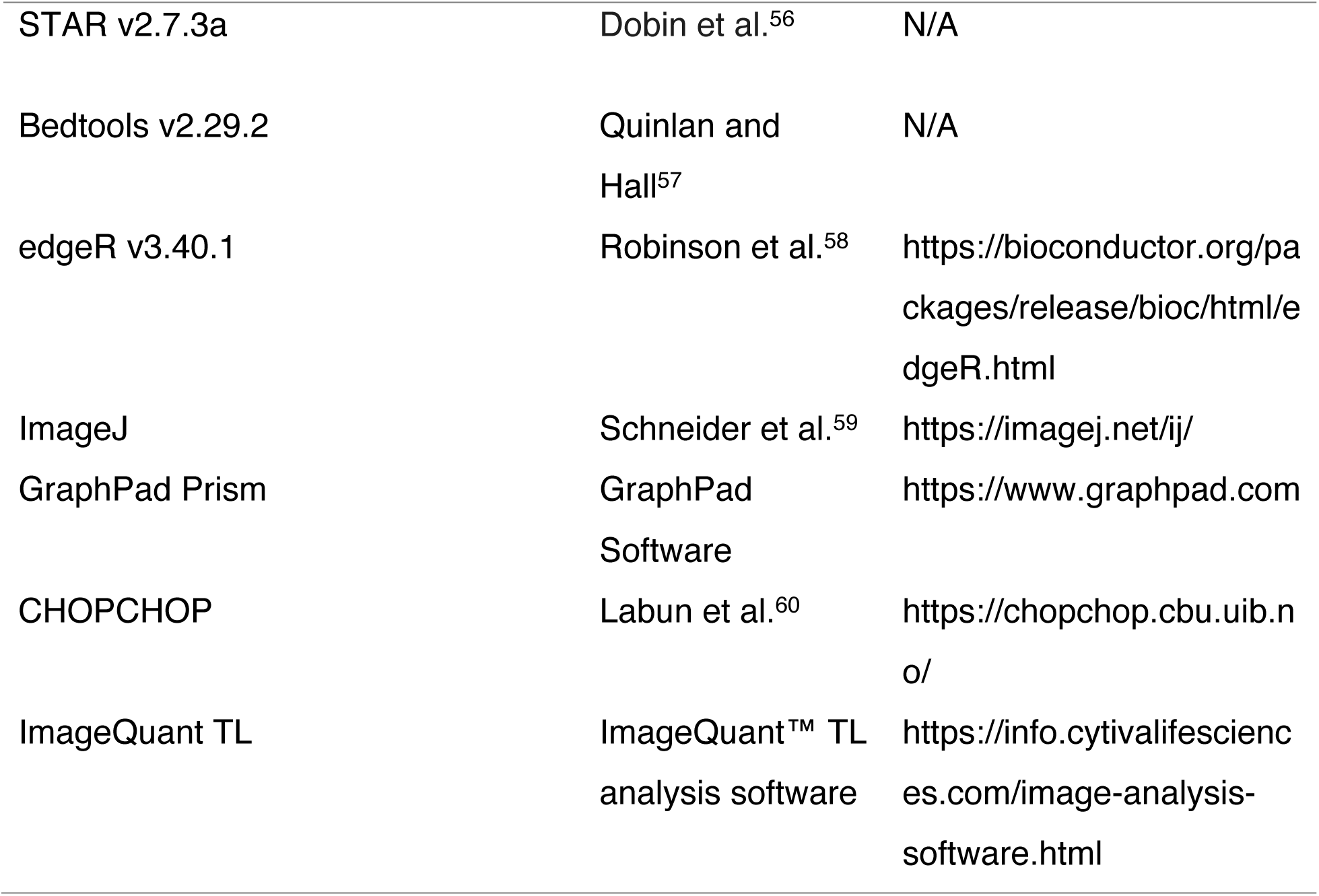

**STAR ★ Methods**
| REAGENT or RESOURCE | SOURCE | IDENTIFIER |
| --- | --- | --- |
| **Antibodies** |  |  |
| Mouse monoclonal anti-FLAG M2 | Sigma-Aldrich | Cat#F1804-200UG; RRID:<br>AB_262044 |
| Rabbit polyclonal anti-beta Tubulin | Invitrogen | Cat#PA1-16947; RRID:<br>AB_795659 |
| Mouse monoclonal anti-GAPDH | Santa Cruz | Cat#sc-365062; RRID:<br>AB_10847862 |
| Rabbit polyclonal anti-CCDC131<br>(ZFC3H1) | Bethyl | Cat#A301-456A; RRID:<br>AB_960958 |
| Rabbit polyclonal anti-MTR4 | Abcam | Cat#AB70551; RRID:<br>AB_1270701 |
| Rabbit polyclonal anti-EXOSC3 | Proteintech | Cat#15062-1-AP; RRID:<br>AB_2278183 |
| Rabbit Polyclonal anti-U2AF35 | Bethyl | Cat# A302-079A; RRID:<br>AB_1604295 |
| Rabbit Polyclonal anti-U2AF65 | Bethyl | Cat#A303-667A; RRID:<br>AB_11204947 |
| Mouse monoclonal anti-U1-70K,<br>clone 9C4.1 | Millipore | Cat#05-1588; RRID:<br>AB_10805959 |
| Mouse monoclonal anti-U1A | Santa Cruz | Cat#sc-101149; RRID:<br>AB_2193721 |
| Rabbit monoclonal anti-FLAG | Cell Signaling | Cat#14793S; RRID:<br>AB_2572291 |
| Rabbit Polyclonal anti-Lamin-A | Bethyl | Cat#A303-433A; RRID:<br>AB_10951693 |
| Mouse monoclonal anti-ENY2 | Invitrogen | Cat# MA5-27843; RRID:<br>AB_2735176 |
| Rabbit Polyclonal anti-Histone H3 | Proteintech | Cat# 17168-1-AP; RRID:<br>AB_2716755 |
| Goat Anti-Rabbit IgG HRP-conjugate | Millipore | Cat#12-348; RRID:<br>AB_390191 |
| Goat Anti-Mouse IgG HRP-conjugate | Millipore | Cat#12-349; RRID:<br>AB_390192 |
| **Chemicals, peptides, and recombinant proteins** |  |  |
| DMEM | Gibco | Cat#11995065 |
| Fetal Bovine Serum | Sigma | Cat#F0926-100ML |
| Lipofectamine 3000 | Invitrogen | Cat#L3000015 |
| In-Fusion Snap Assembly Master Mix | Takara | Cat#638949 |
| Blasticidin | Invivogen | Cat#ant-bl-05 |
| Puromycin | Invivogen | Cat#ant-pr-1 |
| dTAGv-1 | Tocris | Cat#6914 |
| Polybrene | Sigma | Cat#TR-1003 |
| Trizol | Invitrogen | Cat#15596026 |
| NEBNext Magnesium RNA fragmentation module | NEB | Cat#E6150S |
| Phusion High-Fidelity PCR Master Mix with HF Buffer | NEB | Cat#M0531L |
| Lonza SF Cell Line 4D-Nucleofector X Kit | ThermoFisher Scientific | Cat#NC0281111 |
| RQ1 RNase-Free DNase | Promega | Cat#M6101 |
| Superscript III Reverse Transcriptase | ThermoFisher Scientific | Cat#18080044 |
| Random hexamers | ThermoFisher Scientific | Cat#FERSO142 |
| PowerUp™ SYBR™ Green Master Mix | Applied Biosystems | Cat#A25742 |
| Poly-D-Lysine | Gibco | Cat#A3890401 |
| Paraformaldehyde | ThermoFisher Scientific | Cat#AA47377-9L |
| 20X SSC | Invitrogen | Cat#15557-036 |
| Formamide (Deionized) | Ambion | Cat#AM9342 |
| Tween-20 | ThermoFisher Scientific | Cat#AAJ20605AP |
| Dextran Sulfate | Fisher | Cat#BP1585-100 |
| E. coli tRNA | Roche | Cat#10 109 541 001 |
| Vanadyl ribonucleoside complex | NEB | Cat#S1402S |
| DAPI | Invitrogen | Cat#D1306 |
| Fluoromount G | Invitrogen | Cat#00495802 |
| T7 RNA Polymerase | NEB | Cat#M0251S |
| Amersham MicroSpin G-50 Columns | Cytiva | Cat#27533001 |
| Yeast tRNA | Invitrogen | Cat#AM7119 |
| Amylose Resin | NEB | Cat#E8021L |
| Halt™ Protease Inhibitor Cocktail | ThermoFisher Scientific | Cat#78429 |
| ANTI-FLAG M2 Affinity Gel | Millipore | Cat#A2220 |
| FLAG peptide | Sigma-Aldrich | Cat#F3290 |
| RNase A | ThermoFisher Scientific | Cat#EN0531 |
| 0.45 µm nitrocellulose membrane | Bio-Rad | Cat#1620115 |
| **Critical commercial assays** |  |  |
| MycoAlert Mycoplasma Detection Kit | Lonza | Cat#LT07-318 |
| qPCR Lentivirus Titer Kit | Applied Biological Materials | Cat#LV900 |
| 4-20% pre-cast SDS-PAGE gels | BioRad | Cat#4568096 |
| **Deposited data** |  |  |
| mRNA-seq | This study | GEO: GSE299756 |
| PAS-seq | This study | GEO: GSE299756 |
| Proteome | This study | ProteomeXchange:<br>PXD065356 |
| **Experimental models: Cell lines** |  |  |
| HEK293T | ATCC | Cat#CRL-3216 |
| LENG8 dTAG HEK293T | This study | N/A |
| ZFC3H1 dTAG HEK293T | This study | N/A |
| EXOSC3 dTAG HEK293T | This study | N/A |
| ZC3H3 dTAG HEK293T | This study | N/A |
| Flp-in 293-eGFP-5' ss-bGH | This study | N/A |
| Flp-in 293-eGFP-5' ss mut-bGH | This study | N/A |
| **Oligonucleotides** |  |  |
| RT-qPCR primers, shRNA sequences, sgRNA sequence, siRNA sequences (see Table 7) | This paper | N/A |
| Oligod(T)30 | This paper | N/A |
| eGFP FISH probes | Ding and Elowitz<sup>54</sup> | N/A |
| **Recombinant DNA** |  |  |
| LentiCRISPR v2-scramble | This study | N/A |
| LentiCRISPR v2-KHSRP sgRNA-1 | This study | N/A |
| LentiCRISPR v2-KHSRP sgRNA-2 | This study | N/A |
| LentiCRISPR v2-ZNF207 sgRNA-1 | This study | N/A |
| LentiCRISPR v2-ZNF207 sgRNA-2 | This study | N/A |
| LentiCRISPR v2-KIAA1429 sgRNA-1 | This study | N/A |
| LentiCRISPR v2-KIAA1429 sgRNA-2 | This study | N/A |
| LentiCRISPR v2-METTL3 sgRNA-1 | This study | N/A |
| LentiCRISPR v2-METTL3 sgRNA-2 | This study | N/A |
| LentiCRISPR v2-ZC3H13 sgRNA-1 | This study | N/A |
| LentiCRISPR v2-ZC3H13 sgRNA-2 | This study | N/A |
| LentiCRISPR v2-HNRNPD sgRNA-1 | This study | N/A |
| LentiCRISPR v2-HNRNPD sgRNA-2 | This study | N/A |
| LentiCRISPR v2-ZC3H3 sgRNA-1 | This study | N/A |
| LentiCRISPR v2-ZC3H3 sgRNA-2 | This study | N/A |
| LentiCRISPR v2-LENG8 sgRNA-1 | This study | N/A |
| LentiCRISPR v2-LENG8 sgRNA-2 | This study | N/A |
| LentiCRISPR v2-ZFC3H1 sgRNA-1 | This study | N/A |
| LentiCRISPR v2-ZFC3H1 sgRNA-2 | This study | N/A |
| LentiCRISPR v2-TOE1 sgRNA-1 | This study | N/A |
| LentiCRISPR v2-TOE1 sgRNA-2 | This study | N/A |
| LentiCRISPR v2-SNAPC2 sgRNA-1 | This study | N/A |
| LentiCRISPR v2-SNAPC2 sgRNA-2 | This study | N/A |
| LentiCRISPR v2-EXOSC3 sgRNA-1 | This study | N/A |
| LentiCRISPR v2-EXOSC3 sgRNA-2 | This study | N/A |
| LentiCRISPR v2-LUC7L3 sgRNA-1 | This study | N/A |
| LentiCRISPR v2-LUC7L3 sgRNA-2 | This study | N/A |
| LentiCRISPR v2-ZC3H18 sgRNA-1 | This study | N/A |
| LentiCRISPR v2-ZC3H18 sgRNA-2 | This study | N/A |
| pCRIS-PITCHv2-dTAG-BSD<br>(EXOSC3) | This study | N/A |
| pCRIS-PITCHv2-BSD-dTAG<br>(ZFC3H1) | This study | N/A |
| pCRIS-PITCHv2-BSD-dTAG<br>(LENG8) | This study | N/A |
| pCRIS-PITCHv2-BSD-dTAG<br>(ZC3H3) | This study | N/A |
| pCRIS-PITCHv2-dTAG-BSD<br>(ZNF207) | This study | N/A |
| pX330A-S-EXOSC3-sgRNA | This study | N/A |
| pX330A-S-ZFC3H1-sgRNA | This study | N/A |
| pX330A-S-LENG8-sgRNA | This study | N/A |
| pX330A-S-ZC3H3-sgRNA | This study | N/A |
| pX330A-S-ZNF207-sgRNA | This study | N/A |
| pCDNA5: FLAG-eGFP-5'ss WT-bGH PAS | This study | N/A |
| pCDNA5: FLAG-eGFP-5'ss Mut-bGH PAS | This study | N/A |
| pCDNA5: FLAG-eGFP-5'ss WT-MALAT 3' | This study | N/A |
| pCDNA5: FLAG-eGFP-5'ss Mut-MALAT 3' | This study | N/A |
| pCDNA5: FLAG-eGFP-5'ss WT-L3<br>PAS | This study | N/A |
| pCDNA5: FLAG-eGFP-5'ss Mut-L3 PAS | This study | N/A |
| pCDNA5: FLAG-eGFP-5 x BoxB-bGH PAS | This study | N/A |
| pCDNA5: FLAG-eGFP-5 x BoxB-MALAT 3' | This study | N/A |
| pCDNA5: FLAG-eGFP (ATG mut)-5<br>x BoxB-MALAT 3' | This study | N/A |
| pBluescript: 3XMS2-5'ss WT-bGH<br>PAS | This study | N/A |
| pBluescript: 3XMS2-5'ss Mut-bGH<br>PAS | This study | N/A |
| pBluescript: 3XMS2-5'ss WT-L3<br>PAS | This study | N/A |
| pBluescript: 3XMS2-5'ss Mut-L3 PAS | This study | N/A |
| pCDNA3: FLAG-ZFC3H1 Full length | This study | N/A |
| pCDNA3: λN-FLAG-ZFC3H1 Full<br>length | This study | N/A |
| pCDNA3: FLAG-ZNF207 Full length | This study | N/A |
| pCDNA3: λN-FLAG-ZNF207 Full<br>length | This study | N/A |
| pCDNA3: FLAG-ZC3H3 Full length | This study | N/A |
| pCDNA3: λN-FLAG-ZC3H3 Full<br>length | This study | N/A |
| pCDNA3: FLAG-LENG8 Full length | This study | N/A |
| pCDNA3: λN-FLAG-LENG8 Full<br>length | This study | N/A |
| pCDNA3: FLAG-U1A Full length | This study | N/A |
| pCDNA3: λN-FLAG-U1A Full length | This study | N/A |
| pCDNA3: FLAG-U2AF2 Full length | This study | N/A |
| pCDNA3: λN-FLAG-U2AF2 Full<br>length | This study | N/A |
| pCDNA3: FLAG-GANP Full length | This study | N/A |
| pCDNA3: λN-FLAG-GANP Full length | This study | N/A |
| pCDNA3: FLAG-LENG8 SAC3 | This study | N/A |
| pCDNA3: λN-FLAG-LENG8 SAC3 | This study | N/A |
| pCDNA3: FLAG-GANP SAC3 | This study | N/A |
| pCDNA3: λN-FLAG-GANP SAC3 | This study | N/A |
| pCDNA3: FLAG-LENG8 1-540 c-myc NLS | This study | N/A |
| pCDNA3: FLAG-LENG8 541-800 c-myc NLS | This study | N/A |
| pCDNA3: FLAG-LENG8 1-290 c-myc NLS | This study | N/A |
| pCDNA3: FLAG-LENG8 200-400 c-myc NLS | This study | N/A |
| pCDNA3: FLAG-LENG8 290-540 c-myc NLS | This study | N/A |
| pCDNA3: FLAG-LENG8 1-290 mut1 c-myc NLS | This study | N/A |
| pCDNA3: FLAG-LENG8 1-290 mut2 c-myc NLS | This study | N/A |
| pCDNA3: FLAG-LENG8 1-290 N36A c-myc NLS | This study | N/A |
| pCDNA3: FLAG-LENG8 1-290 E38A/W39A c-myc NLS | This study | N/A |
| pCDNA3: FLAG-LENG8 1-290 Q82A c-myc NLS | This study | N/A |
| pCDNA3: FLAG-LENG8 1-290 Y86A/Y87A c-myc NLS | This study | N/A |
| pCDNA3: FLAG-LENG8 1-290<br>W89A/Y90A c-myc NLS | This study | N/A |
| pCDNA3: FLAG-LENG8 1-290<br>Y93A/N94A c-myc NLS | This study | N/A |
| pCDNA3: FLAG-LENG8 1-290<br>Y97A/Y98A c-myc NLS | This study | N/A |
| pCDNA3: FLAG-LENG8 200-400<br>mut5 c-myc NLS | This study | N/A |
| pCDNA3: FLAG-LENG8 200-400<br>E307A/K310A c-myc NLS | This study | N/A |
| pCDNA3: FLAG-LENG8 200-400<br>D311A/E314A c-myc NLS | This study | N/A |
| pCDNA3: FLAG-LENG8 200-400<br>mut5 c-myc NLS | This study | N/A |
| pCDNA3: FLAG-LENG8 Full length<br>R563A | This study | N/A |
| pCDNA3: FLAG-LENG8 Full length<br>Mut5 | This study | N/A |
| pCDNA3: FLAG-LENG8 Full length<br>E38A/W39A | This study | N/A |
| pCDNA3: FLAG-Chimera | This study | N/A |
| pCDNA3: λN-FLAG-Chimera | This study | N/A |
| pCDNA3: Strep-ZFC3H1 600-990 c-myc NLS | This study | N/A |
| pCDNA3: Strep-ZFC3H1 600-990<br>E735A/R742A/E746A c-myc NLS | This study | N/A |
| **Software and algorithms** |  |  |
| Cutadapt v2.10 | Martin<sup>55</sup> | N/A |
| STAR v2.7.3a | Dobin et al.<sup>56</sup> | N/A |
| Bedtools v2.29.2 | Quinlan and Hall<sup>57</sup> | N/A |
| edgeR v3.40.1 | Robinson et al.<sup>58</sup> | https://bioconductor.org/packages/release/bioc/html/edgeR.html |
| ImageJ | Schneider et al.<sup>59</sup> | https://imagej.net/ij/ |
| GraphPad Prism | GraphPad Software | https://www.graphpad.com |
| CHOPCHOP | Labun et al.<sup>60</sup> | https://chopchop.cbu.uib.no/ |
| ImageQuant TL | ImageQuant™ TL analysis software | https://info.cytivalifesciences.com/image-analysis-software.html |

### Experimental model and study participant details

#### Cell Lines

All HEK293T cell lines were cultured in DMEM (Gibco, 11995065) supplemented with 10% FBS (Sigma, F0926-100ML). Flp-In T-REx HEK293 cells were maintained in the same medium with the addition of 100 μg/mL Zeocin (Invivogen, NC9002627) and 15 μg/mL blasticidin (Invivogen, ant-bl-1). For Flp-In T-REx HEK293 cell lines harboring either eGFP-5′ ss-PAS or eGFP-5′ ss mut-PAS reporters, the medium was further supplemented with 15 μg/mL blasticidin and 500 μg/mL G418 (Corning, MT30234CR). All cells were incubated at 37 °C with 5% CO₂ and routinely tested for mycoplasma contamination using the MycoAlert Mycoplasma Detection Kit (Lonza, LT07-318).

### Method details

#### Generation of stable Flp-In T-REx HEK293 Cell Lines Expressing eGFP-5′ ss-PAS or eGFP-5′ ss mut-PAS Reporters

To generate stable Flp-In T-REx HEK293 cell lines expressing either the eGFP-5′ ss-PAS or eGFP-5′ ss mut-PAS reporter, we followed the manufacturer’s protocol (Invitrogen, R780-07). Briefly, Flp-In T-REx HEK293 cells were co-transfected with pOG44 and pcDNA5/FRT/TO plasmids at a 9:1 ratio, with the latter encoding either the eGFP-5′ ss-PAS or eGFP-5′ ss mut-PAS reporter. Transfections were carried out using Lipofectamine 3000 (Invitrogen, L3000015). Forty-eight hours post-transfection, cells were split and subjected to dual selection with blasticidin (Invivogen, NC9016621) and G418 (Corning, MT30234CR). The selection medium was refreshed every 2 days to facilitate polyclonal selection of isogenic populations. Individual colonies were subsequently isolated and expanded into single-clone cell lines.

#### Targeted Knock-In of FLAG-FKBP12^F36V^ at the N- or C-Terminus of Endogenous ZFC3H1, ZNF207, LENG8, and ZC3H3 Genes in HEK293T Cell**s**

C-terminal dTAG-FLAG knock-in of ZNF207, and N-terminal FLAG-dTAG knock-in of ZFC3H1, LENG8, and ZC3H3 in HEK293T cells were generated using CRISPR/Cas9 in combination with the microhomology-mediated end joining (MMEJ) strategy, as previously described^30,31^. Donor vectors were constructed by modifying pCRIS-PITChv2-dTAG-BSD (BRD4) (Addgene, #91795) and pCRIS-PITChv2-BSD-dTAG (BRD4) (Addgene, #91792) to replace the HA tag with a 3×FLAG tag. Subsequently, gene-specific microhomology arms for ZFC3H1, ZNF207, LENG8, and ZC3H3 were introduced via In-Fusion cloning (Takara). Plasmids pCRIS-PITChv2-dTAG-BSD (BRD4) and pCRIS-PITChv2-BSD-dTAG (BRD4) were gifts from James Bradner and Behnam Nabet. Guide RNAs were designed using the CHOPCHOP online tool to ensure specificity and efficiency^60^. Single-cell clones were screened and validated by genomic DNA PCR, followed by sanger sequencing when possible, and confirmed by western blotting for all lines. Homozygous knock-in clones were selected for downstream experiments. Notably, for ZFC3H1 and ZC3H3, only the longest isoform showed dTAG-dependent degradation.

#### CRISPR screen

The Human CRISPR Knockout Pooled Library (Brunello) targeting 19,114 genes with a total of 77,441 sgRNAs (4 sgRNAs per gene) was obtained from Addgene^26^ (Pooled Library #73179). Lentiviral particles were generated by co-transfecting pooled library plasmids with packaging plasmids (psPAX2 and pMD2.G) into HEK293T cells at a 4:3:2 ratio using Lipofectamine 3000 (Invitrogene, Cat # L3000015). After 24 hours of incubation, the culture medium was replaced with fresh DMEM containing 10% FBS. Lentiviral supernatants were collected at 48- and 72-hours post-transfection, followed by centrifugation (2000 rpm, 5 min) to remove cell debris. The supernatants were then aliquoted and stored at −80°C until further use. Before infecting reporter cell lines, the virus titer test was performed by qPCR Lentivirus Titer Kit (Applied Biological Materials Inc, Cat. No. LV900)

For CRISPR screens, eGFP-5′ ss-PAS reporter cell lines were infected with the Brunello genome-wide library at a low multiplicity of infection (MOI ∼0.3). After 24 hours, the culture medium was replaced with fresh DMEM containing 0.75 µg/ml puromycin.

Following a 5-day selection period, doxycycline was added to the culture medium at a concentration of 1 µg/µl to induce eGFP expression. After three days of induction, the cells were collected for sorting based on high eGFP expression.

For sorting high eGFP-expressing cell lines, sgRNA-infected cells were resuspended in FACS buffer (1x PBS containing 2% FBS and 0.5M EDTA) to ensure a cell concentration of approximately 10⁶ to 10⁷ cells/ml. The cell suspension was then filtered using a 40 µm nylon filter (Thermo Fisher) before sorting. Based on the controls, the top 1% of cells with the highest eGFP expression were sorted into a collection buffer (50% PBS + 50% FBS) for DNA preparation.

Genomic DNA was extracted from the sorted cells using the Qiagen Blood & Cell Culture DNA Mini Kit. A two-step PCR reaction was then performed to clone and amplify the sgRNA barcode. For the first PCR, the following primers were used: Forward: tatcttgtggaaaggacga; Reverse: gagccaattcccactcctttc. The PCR reaction was prepared using Ex Taq DNA polymerase (Takara, RR001A) and carried out for 18 cycles under the following conditions: 95°C for 30 seconds, 53°C for 30 seconds, and 72°C for 30 seconds. For the second PCR, P5 and P7 sequencing primers were used according to the protocol available on the Addgene website (https://media.addgene.org/cms/filer_public/61/16/611619f4-0926-4a07-b5c7-e286a8ecf7f5/broadgpp-sequencing-protocol.pdf). The PCR reaction was carried out for 14 cycles under the following conditions: 95°C for 30 seconds, 53°C for 30 seconds, and 72°C for 30 seconds.Following amplification, the PCR samples were loaded onto a 2% agarose gel and run at 120V for approximately 35 minutes. The DNA bands were then extracted and purified using AMPure XP-PCR purification for next-generation sequencing (NGS). The input library and 3 replicates of CRISPR screen samples were sequenced by Illumina sequencing. sgRNA sequences were obtained by removing the adaptor sequences and quantified. The gene enrichment levels and false discover rate were obtained using the MAGeCK package.

#### In Vitro RNA Pull-Down Assay

A total of 30 pmol of in vitro-transcribed RNA and 225 pmol of MBP-MS2 fusion protein were combined in Buffer D-100 (20 mM HEPES-NaOH pH 7.9, 100 mM NaCl, 1 mM MgCl₂, 0.2 mM EDTA) in a final volume of 50 µL and incubated on ice for 30 minutes to allow complex formation. Subsequently, 10 µL of 0.1 M ATP, 20 µL of 1 M creatine phosphate, 10 µL of 10 µg/µL yeast tRNA, and 400 µL of HeLa nuclear extract were added, and the reaction was brought to a final volume of 1000 µL with nuclease-free water. The mixture was incubated at 30°C for 20 minutes to allow assembly of ribonucleoprotein complexes. Following incubation, the reaction was chilled on ice and centrifuged at 14,000 rpm (≥16,000 × g) for 1 minute at 4°C to remove insoluble material. The clarified supernatant was transferred to 140 µL of pre-washed amylose resin slurry and incubated with rotation at 4°C for 1 hour. Beads were subsequently washed three times with 1 mL of wash buffer (20 mM HEPES-NaOH pH 7.9, 100 mM KCl, 4% glycerol, 1 mM DTT) supplemented with 0.05% IGEPAL CA-630, followed by one additional wash with detergent-free wash buffer. Bound proteins were eluted twice by incubating the beads with 200 µL of wash buffer (without detergent) supplemented with 20 mM maltose for 10 minutes per elution. Eluates were pooled and subjected to acetone precipitation overnight at −20°C. The next day, samples were centrifuged at 14,000 rpm for 20 minutes at 4°C. Pellets were resuspended in 50 µL of digestion buffer (8 M urea, 25 mM ammonium bicarbonate) for mass spectrometry analysis.

#### FLAG-Immunoprecipitation

HEK293T cells were seeded in 10-cm dishes and transfected the following day with pcDNA3 expression plasmids encoding either full-length or truncated versions of LENG8 or GANP. Forty-eight hours post-transfection, cells were harvested by scraping into 5 mL of cold 1× PBS, followed by centrifugation at 800 × rpm for 5 minutes. Cell pellets were lysed in 1 mL of cold lysis buffer per dish (20 mM HEPES-NaOH pH 7.9, 150 mM NaCl, 0.2% NP-40, and 1× Thermo Scientific Halt™ Protease Inhibitor Cocktail, added fresh). Lysates were sonicated on ice using a microtip sonicator (amplitude 1; 3 seconds on, 10 seconds off; six cycles total). Sonicated lysates were cleared by centrifugation at 14,000 × rpm for 20 minutes at 4 °C, and an aliquot of the supernatant was reserved as input. For immunoprecipitation, 40 µL of Anti-FLAG M2 agarose bead slurry (Sigma) per sample was washed three times with 1 mL of lysis buffer. For RNase-treated conditions, 10 µL of RNase A/T1 was added to the cleared lysate and mixed thoroughly. Supernatants were then incubated with pre-washed FLAG beads at 4 °C for 2 hours with gentle rotation. Beads were washed five times with 1 mL of lysis buffer (10 minutes per wash), and bound proteins were eluted three times using 150 µL of lysis buffer supplemented with 3× FLAG peptide (10 minutes per elution). Combined eluates were acetone-precipitated overnight at –20 °C. The following day, samples were centrifuged at 14,000 × rpm for 20 minutes. Resulting protein pellets were resuspended either in 50 µL of digestion buffer (8 M urea, 25 mM ammonium bicarbonate) for mass spectrometry analysis, or in 30 µL of 1× SDS loading buffer and denatured at 95 °C for 10 minutes for western blot analysis.

#### Mass Spectrometry Sample Preparation and Analysis

Protein samples in 50 µL of digestion buffer were loaded onto pre-rinsed Centricon-10 kDa centrifugal filters (Millipore, UFC501008). Filters were first conditioned with 200 µL of HPLC-grade water and centrifuged at 14,000 × g for 10 minutes at room temperature (RT). After loading the samples, the filters were centrifuged at 14,000 × g for 20 minutes at RT. During this step, a 15 mM TCEP solution was prepared by mixing 0.9 µL of 1 M TCEP with 59.1 µL of digestion buffer. A total of 60 µL of the TCEP solution was added directly to the filter unit, and the samples were incubated at RT for 30 minutes to allow disulfide bond reduction. Meanwhile, a 0.5 M iodoacetamide solution was prepared by dissolving 0.185 g of iodoacetamide in 2 mL of HPLC-grade water. Following reduction, iodoacetamide was added to the sample to a final concentration of 45 mM (5.9 µL into 60 µL of the TCEP solution), and the reaction was incubated in the dark at RT for 30 minutes. The sample was then centrifuged at 14,000 × g for 5–10 minutes to remove excess reagents, washed with 60 µL of digestion buffer, and centrifuged again at 14,000 × g for 20 minutes. The flow-through was discarded. For enzymatic digestion, 60 µL of digestion buffer containing LysC (NEB, P8109S) was added to the filter. The amount of LysC was adjusted to 1% of the estimated total protein content (e.g., 1 µg of LysC for 100 µg of protein), with the enzyme reconstituted in its provided buffer. Samples were incubated at 37°C for 4 hours. Following LysC digestion, the urea concentration was diluted to 1–1.5 M by adding 25 mM ammonium bicarbonate, and trypsin (Promega, V5113) was added at 2% of the total protein content (e.g., 2 µg for 100 µg protein), also reconstituted in its respective buffer. Trypsin digestion was carried out overnight at 37°C with intermittent mixing. The following day, digested peptides were recovered by centrifugation at 14,000 × g for 25 minutes into a clean collection tube. To stop digestion and acidify the eluate for downstream processing, trifluoroacetic acid (TFA) was added to a final concentration of 0.1% using a 1% TFA stock solution. Finally, the peptides were desalted and cleaned using Sep-PAK C18 cartridges (Waters, WAT054955) according to the manufacturer’s instructions. After the final elution, the collected peptides were concentrated using vacuum centrifugation for mass spectrometry analysis. Mass spectrometry analysis was performed using Thermo Scientific Orbitrap Fusion Lumos Tribrid Mass Spectrometer at UC Irvine High-end Mass Spectrometry Facility. The change in percent coverage of each protein in different groups were calculated and ranked.

#### PAS-seq analysis

PAS-seq libraries were prepared as previously described with minor modifications^37^. Purified mRNA was fragmented at 94°C for 3 minutes using the NEBNext Magnesium RNA Fragmentation Module (NEB). Libraries were resolved on a 2.5% agarose gel, and fragments in the 200–300 base pair range were excised and purified by gel extraction prior to sequencing on the Illumina NovaSeq 6000 platform. PAS-seq data processing was performed as previously described. Briefly, raw reads were trimmed using Cutadapt (v2.10) to remove (1) a 6-nucleotide linker sequence, (2) poly(A) tails, and (3) any reads that failed trimming^55^. Cleaned reads were then aligned to a concatenated human (hg19) reference genome using STAR (v2.7.3a) with the alignEndsType EndToEnd parameter to ensure full-length alignment^56^. The resulting BAM files were converted to BED format using BEDTools (v2.29.2)^57^. To remove potential internal priming artifacts, a custom Python script was used to filter out reads mapping to genomic regions with evidence of A-rich sequences—specifically, regions containing either six consecutive adenosines or seven adenosines within a 10-nucleotide window downstream of the read. The filtered reads were converted back to BAM format for downstream analysis, including the generation of BigWig coverage tracks using BEDTools. To identify and quantify polyadenylation sites, the 3′ end positions of the filtered reads were extracted using bedtools flank and read counts for each poly(A) site were calculated with bedtools coverage against a master annotation file of known polyadenylation sites. To detect differentially expressed transcripts using PAS-Seq data, we treated counts mapping to each poly(A) site region as a “gene” and analyzed differential gene expression using the edgeR (v3.40.1)^58^. We considered a transcript to be differentially expressed if it displayed an FDR ≤ 0.05 and a log2FC > 1 or log2FC < -1, unless indicated otherwise.

#### mRNA-seq analysis

Total RNAs were extracted and poly(A) RNAs were isolated and the mRNA-seq libraries were prepared by the UCI Genomics Research and Technology Hub (GRTH) and subjected to Illumina Sequencing. The sequencing reads were filtered for quality, adaptors were removed by using Cutadapt^55^, and the reads were mapped to the human genome (hg19) using STAR^56^. Intron retention analysis was performed by using rMATS^61^.

#### RNA FISH and Immunofluorescence

To assess RNA localization of eGFP reporters, HEK293T cells were seeded in 8-well glass-bottom chamber slides (ibidi) pre-coated with 0.1 mg/mL Poly-D-Lysine (Gibco). The following day, cells were transfected with 100 ng of each eGFP reporters using Lipofectamine 3000 (Invitrogen). Twenty-four hours post-transfection, cells were fixed with 4% paraformaldehyde for 10 minutes, washed with 1× PBS, and permeabilized in 70% ethanol at –20 °C overnight. RNA-FISH probes targeting the eGFP coding sequence were synthesized as previously described. For hybridization, cells were washed once with wash buffer (2× SSC, 20% deionized formamide, 0.1% Tween-20) and incubated overnight at 30 °C in 100 µL of hybridization buffer containing 125 ng of RNA-FISH probes. The hybridization buffer included 0.1 g/mL dextran sulfate (Fisher), 0.1 mg/mL E. coli tRNA (Roche), 2 mM vanadyl ribonucleoside complex (NEB), 2× SSC (Invitrogen), 20% deionized formamide (Ambion), and 0.1% Tween-20. The following day, cells were washed twice with pre-warmed wash buffer at 30 °C, followed by a 30-minute incubation in fresh wash buffer. Nuclei were then stained with DAPI (Invitrogen, 1:20,000 dilution) for 5 minutes and mounted with Fluoromount-G (Invitrogen) prior to imaging.

For combined RNA FISH and immunofluorescence, cells were post-fixed with 4% paraformaldehyde for 10 minutes after hybridization. Immunostaining was performed by incubating cells with primary antibodies for 2 hours at room temperature in PBS containing 0.1% Tween-20 (PBST), followed by a single PBST wash and 1 hour incubation with fluorescent secondary antibodies. After additional PBST washes, nuclei were stained with DAPI for 5 minutes, and cells were mounted with Fluoromount-G prior to imaging. Images were acquired using a Zeiss LSM 900 Airyscan 2 microscope.

#### RT-qPCR analysis

Total RNA was extracted from cells using TRIzol reagent (Invitrogen, 15596018) according to the manufacturer’s instructions. Reverse transcription was performed using All-In-One 5X RT MasterMix with gDNA Removal (Applied Biological Materials, G592), following the provided protocol. The resulting cDNA was used for quantitative PCR (qPCR) with PowerUp™ SYBR™ Green Master Mix (Applied Biosystems, A25742). The relative abundance of transcripts was quantified using the ΔΔCt method.

#### Western blot analysis

Cells were lysed in RIPA buffer (50 mM Tris-HCl, pH 8.0; 150 mM NaCl; 1% NP-40; 0.5% sodium deoxycholate; 0.1% SDS), and lysates were mixed with SDS loading dye. Samples were denatured at 95 °C for 10 minutes, resolved on a 10% polyacrylamide gel, and transferred onto a 0.45 µm nitrocellulose membrane (Bio-Rad, 1620115) using the eBlot L1 wet-transfer system (GenScript, L00686) with the standard program. Following transfer, membranes were blocked in 5% non-fat milk prepared in PBST (0.1% Tween-20 in PBS) for 1 hour at room temperature, then incubated overnight at 4 °C with primary antibodies diluted in 5% milk in PBST. The next day, membranes were washed with PBST and incubated with HRP-conjugated secondary antibodies in 5% milk in PBST for 1 hour at room temperature. After final washes in PBST, signals were developed using Radiance Q Chemiluminescent Substrate (Azure Biosystems, 10147-296) and imaged with a Bio-Rad ChemiDoc MP system.

#### U1 snRNP Inhibition by Antisense Morpholino Oligonucleotides (AMOs)

To inhibit U1 snRNP, 1 × 10⁶ cells were nucleofected with 25 µM control or U1 antisense morpholino oligonucleotides (AMOs; GeneTools, LLC) using the Lonza SF Cell Line 4D-Nucleofector™ X Kit and the DH-135 program. Cells were imaged 8 hours post-nucleofection using a fluorescence microscope.

#### Construction of eGFP Reporters

The eGFP-5′ ss-PAS reporter is consisted of a FLAG-eGFP coding sequence followed by a 28-nucleotide 5′ ss sequence derived from the NXF1 gene, previously shown to include both a strong and a weaker 5′ ss^25^. In the eGFP-5′ ss mut-PAS reporter, both splice sites were disrupted via PCR-based mutagenesis. Polyadenylated versions of these reporters contained either the bovine growth hormone (bGH) polyadenylation signal or the adenovirus major late (L3) poly(A) site downstream of the 5′ ss. In contrast, non-polyadenylated reporters incorporated a 164-nucleotide sequence from the 3′ end of MALAT1^35^.

The eGFP-BoxB reporter consists of a FLAG-eGFP coding sequence followed by five tandem BoxB motifs (5xBoxB), which serve as binding sites for the λN peptide, enabling targeted tethering of specific proteins to the reporter RNA^36^. In polyadenylated versions, bGH polyadenylation signal was placed downstream of the 5xBoxB sequence. In contrast, non-polyadenylated versions incorporated a 164-nucleotide sequence from the 3′ end of MALAT1.

#### RNA Pulldown Constructs

The 5′ ss RNA pulldown constructs were cloned into the pBluescript vector and contained three tandem MS2 hairpin sequences followed by either a 28-nucleotide 5′ ss sequence derived from the NXF1 gene or a mutant 5′ ss motif, along with a bGH poly(A) signal.

#### Generation of FLAG-Tagged Expression Constructs

Expression constructs for FLAG-LENG8, FLAG-ZFC3H1, FLAG-ZNF207, and FLAG-ZC3H3 and Flag-GANP were cloned into the pCDNA3 vector. For the tethering assay, the λN-peptide was inserted at the N-terminus of each protein, immediately downstream of the FLAG tag. Truncation and point mutation variants were generated via PCR-based mutagenesis. All truncated constructs included a C-terminal c-Myc nuclear localization signal (NLS).

#### Lentiviral knockout

Lentiviral particles were produced by co-transfecting HEK293T cells with the plasmids lentiCRISPRv2, PMD2.G, and psPAX2. Twenty-four hours post-transfection, the medium was replaced with fresh medium of the same composition. Virus-containing supernatant was collected at 48- and 72-hours post-transfection and filtered through a 0.45 µm filter. Viral titers were quantified using a qPCR-based kit (Abm Good). HEK293T cells were transduced at a multiplicity of infection (MOI) of 10 with lentivirus encoding either sgRNAs targeting candidate factors or a non-targeting scramble control. Polybrene (8 µg/mL) was added to enhance transduction efficiency. Twenty-four hours after transduction, puromycin was added to the culture medium at a final concentration of 1.25 µg/mL. Following a 5-day selection period, doxycycline was added at 1 µg/µL to induce eGFP expression. Three days after induction, eGFP intensity was analyzed by flow cytometry (FACS).

#### Subcellular Fractionation for Protein and RNA Isolation

When cell confluency reached 90%, the culture medium was discarded, and the cells were washed three times with pre-chilled 1× PBS. After completely removing the PBS, trypsin was added to digest the cells. Upon completion of digestion, complete culture medium was added to neutralize the trypsin. Cells were gently pipetted to form a uniform suspension and transferred to a 15 mL centrifuge tube. The suspension was centrifuged at 800 rpm for 5 minutes, and the supernatant was discarded. The pellet was resuspended in 1 mL of 1× PBS, transferred to a 1.5 mL microcentrifuge tube, and centrifuged again at 800 rpm for 5 minutes. The supernatant was carefully removed. To lyse the cells, 150 μL of pre-chilled 0.7% NP-40 lysis buffer (10 mM Tris-HCl, 75 mM NaCl, 0.7% NP-40, pH 7.5, supplemented with fresh protease inhibitors) was added. The suspension was pipetted using a 200 μL tip until homogeneous and incubated on ice for 30 seconds. Samples were then centrifuged at 13,300 rpm for 15 seconds to separate the cytoplasmic (supernatant) and nuclear (pellet) fractions. A total of 150 μL of the cytoplasmic supernatant was collected into new tubes for western blotting or cytoplasmic RNA extraction by TRIzol reagent. The nuclear pellet was washed three times with 1× PBS. For downstream applications, either 100 μL of RIPA buffer (with fresh protease inhibitors) or 1 mL of TRIzol reagent was added to extract nuclear proteins or nuclear RNAs, respectively.

---

**S1.**
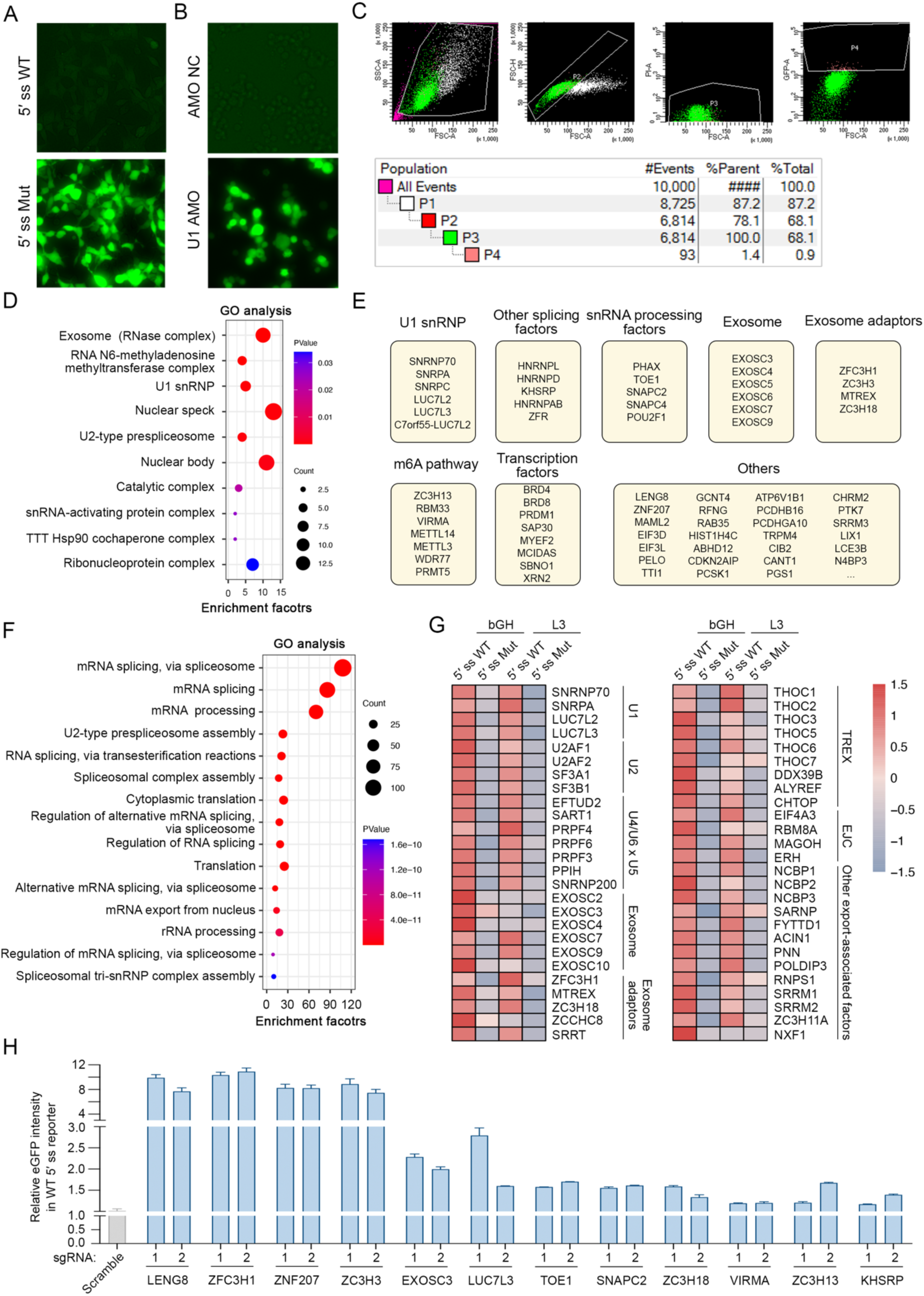

**S2.**
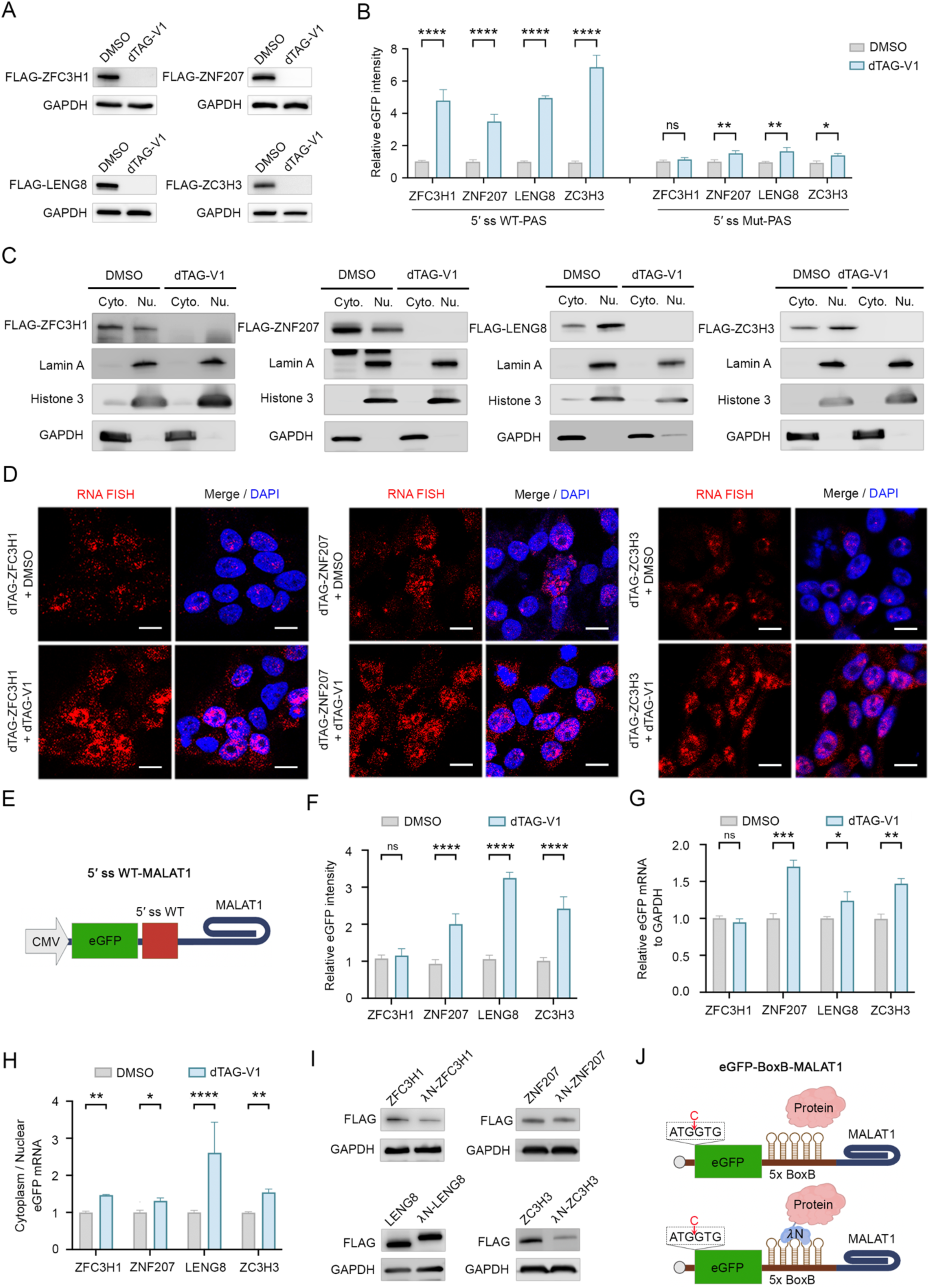

**S3.**
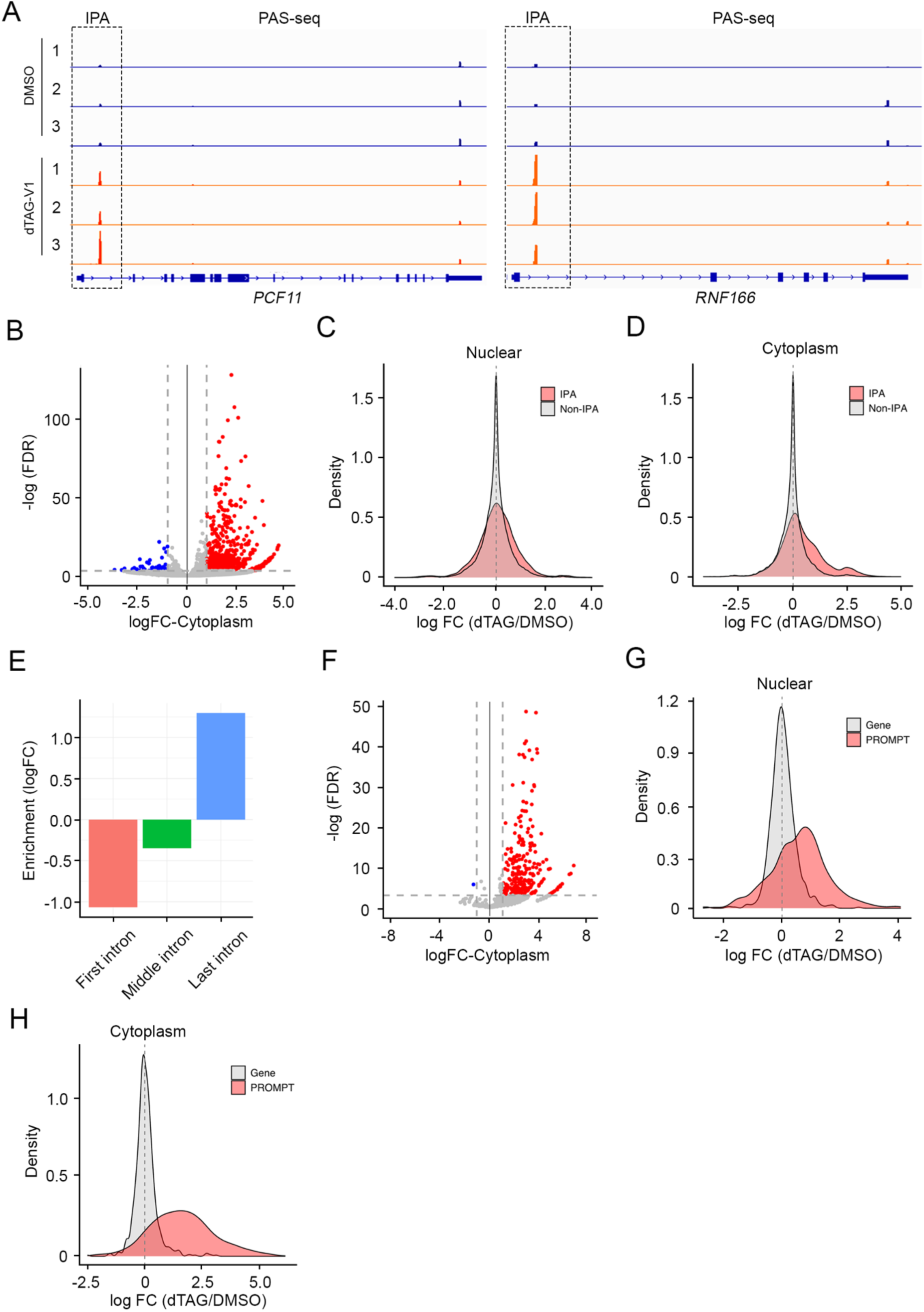

**S4.**
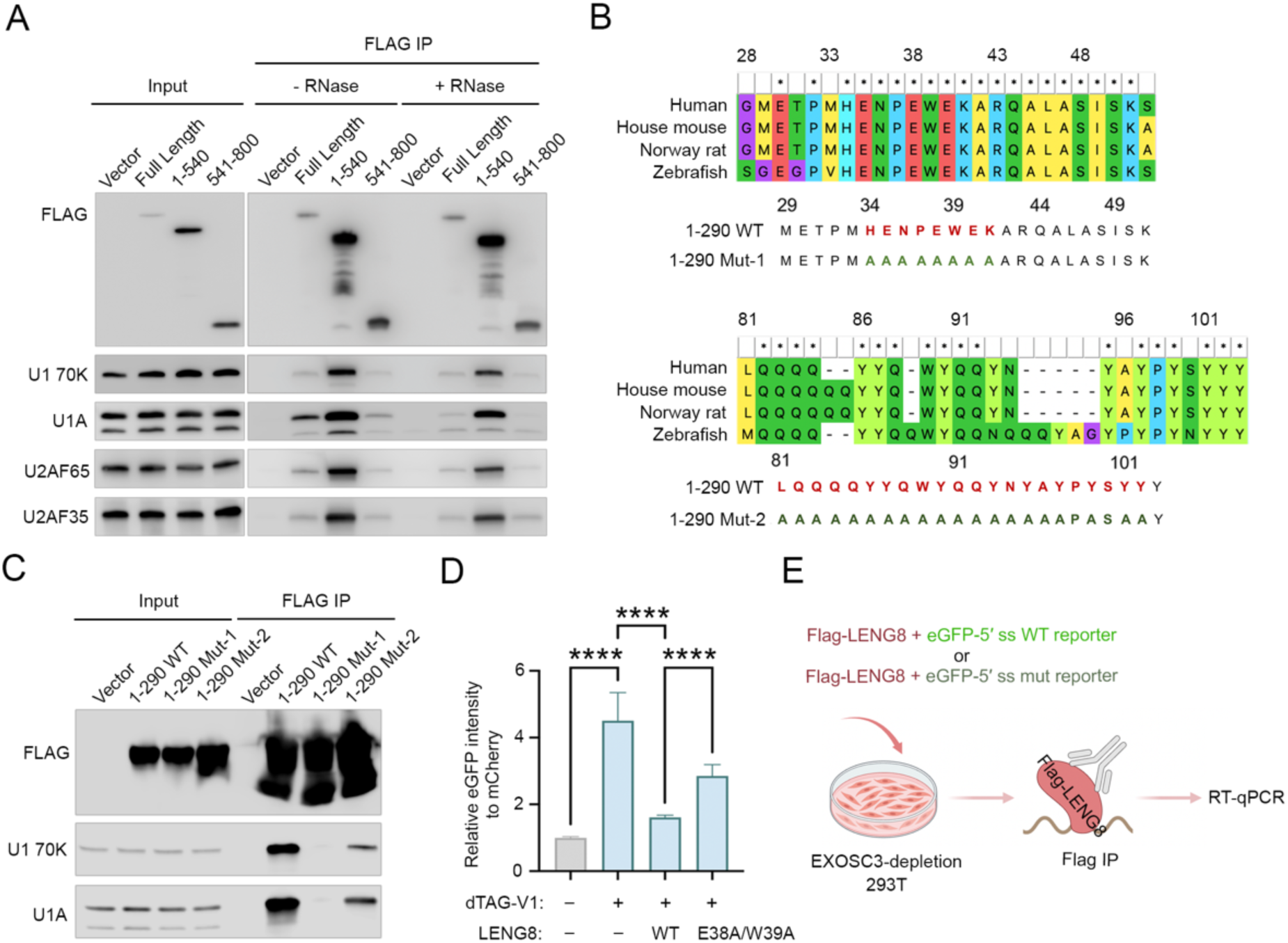

**S5.**
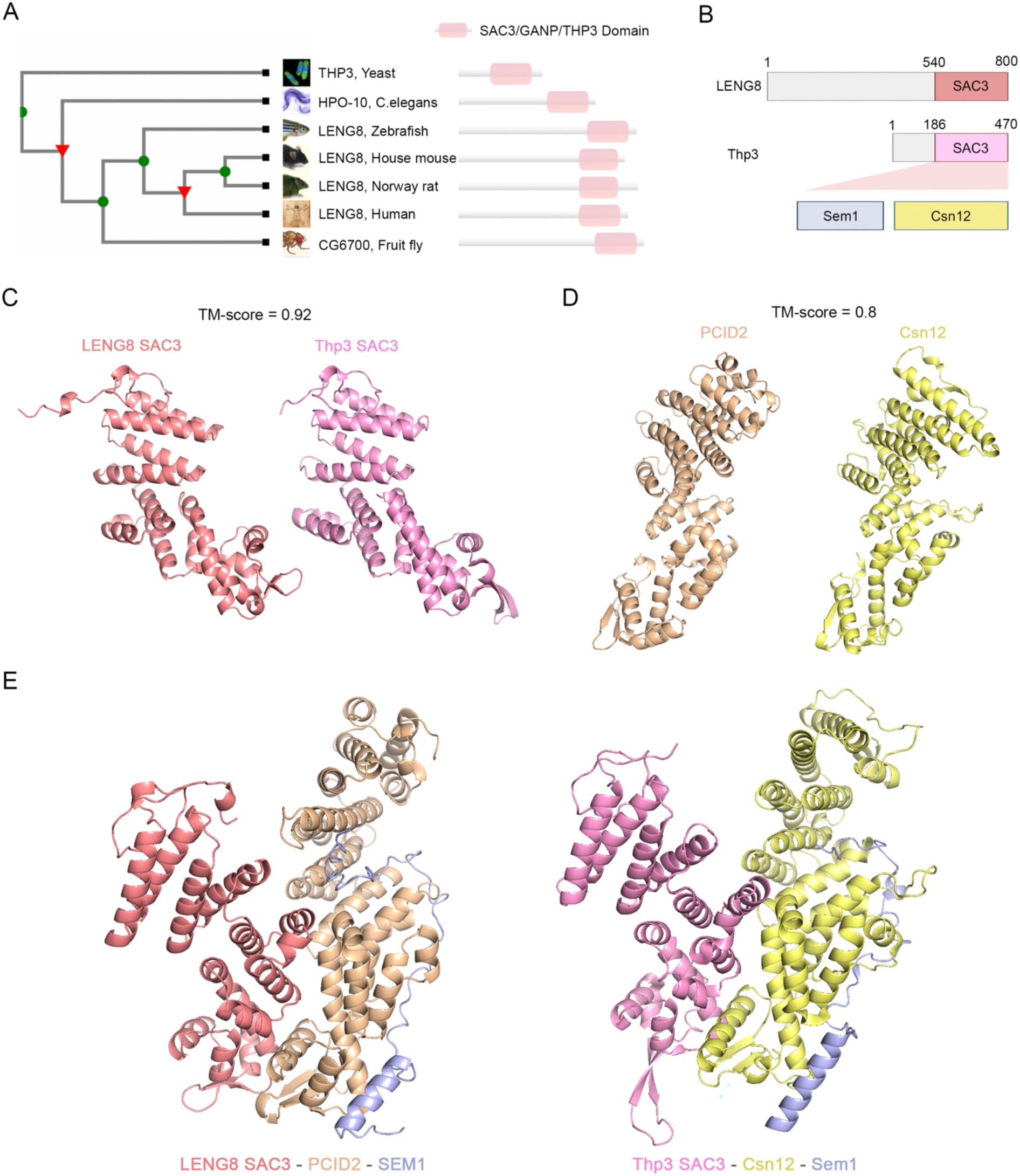

**S6.**
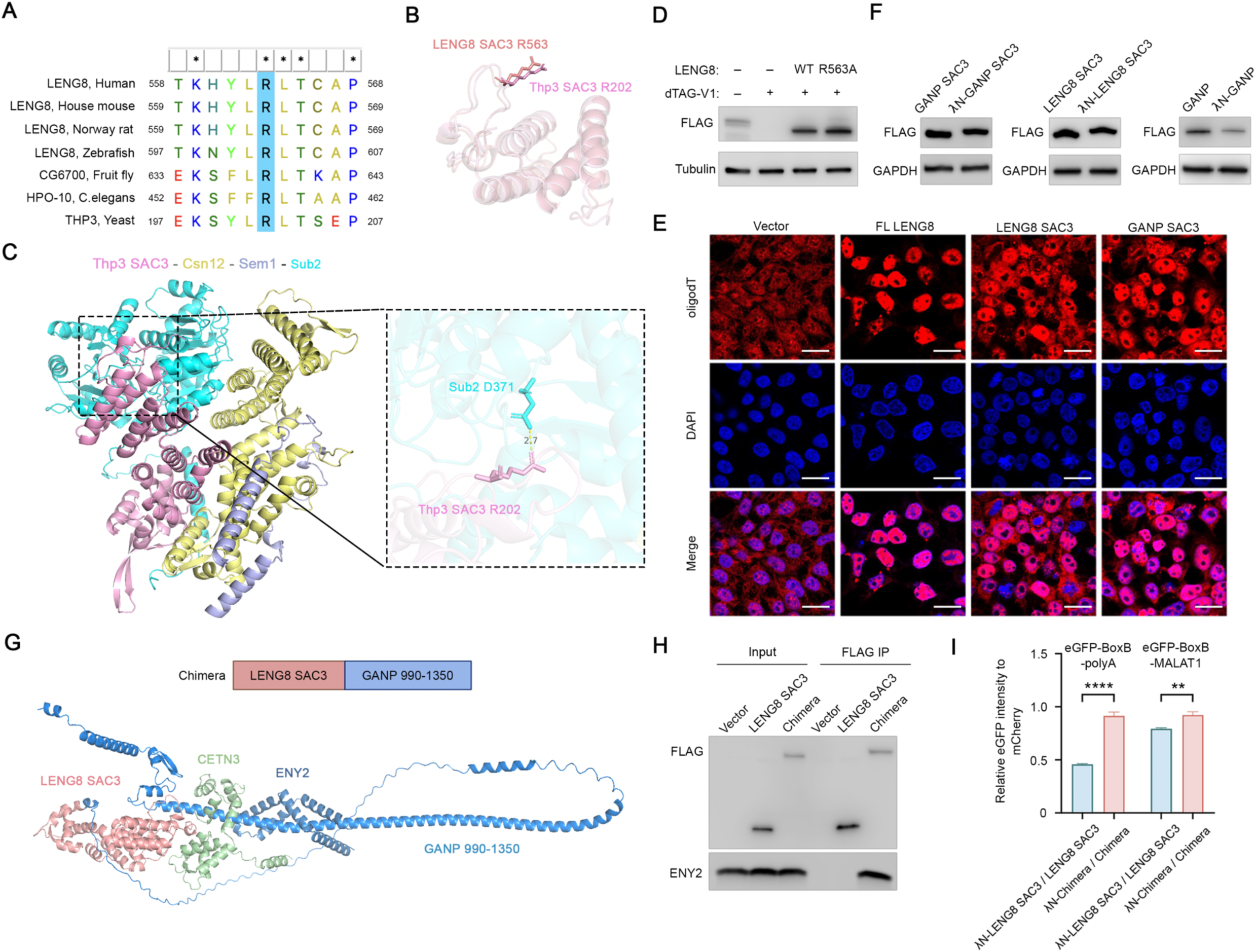

**S7.**
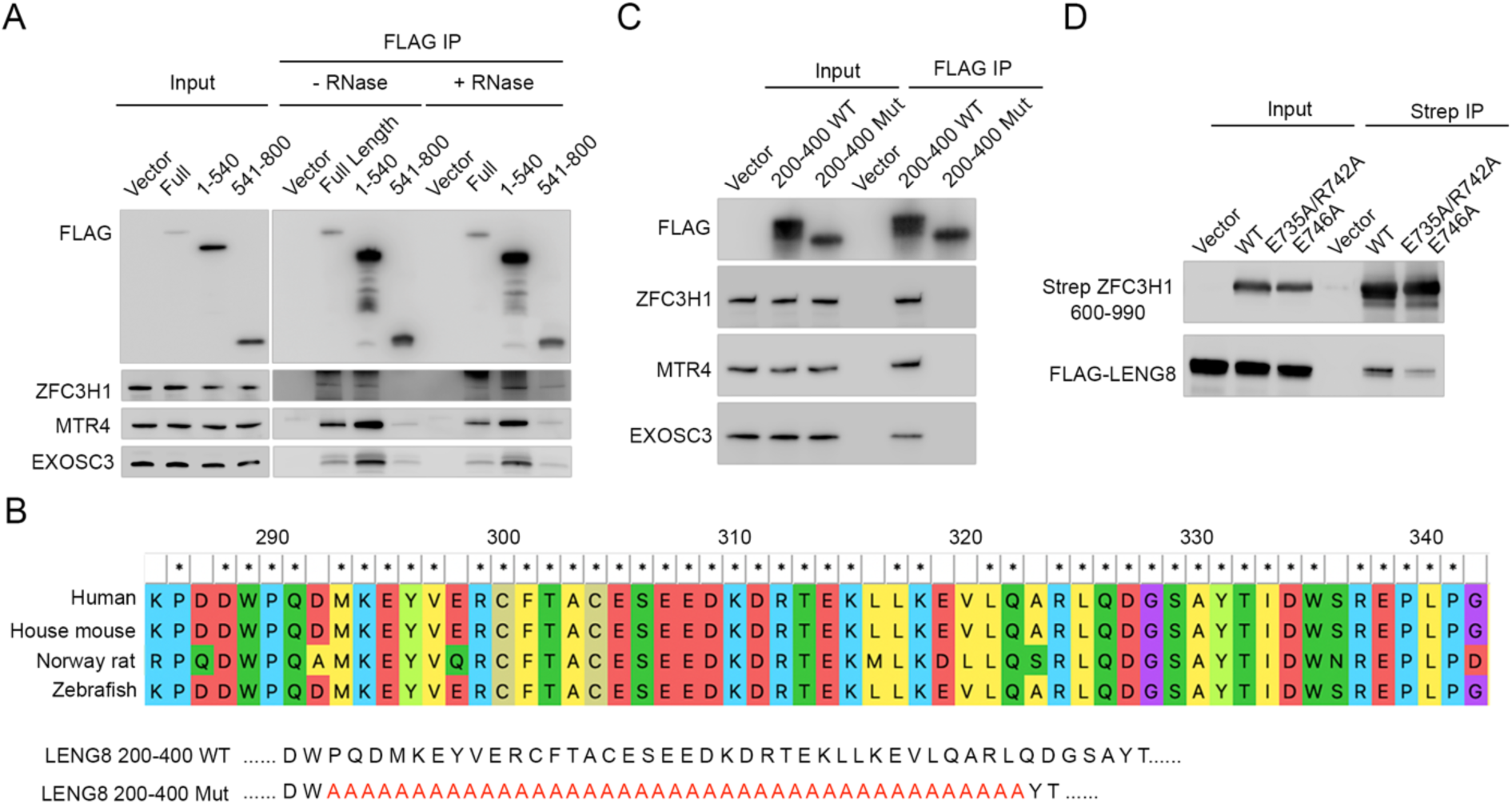

